# Matrix-controlled emergence of biofilm architecture shapes antimicrobial survival

**DOI:** 10.64898/2026.09.25.752283

**Authors:** Tom E. R. Belpaire, Jolien J. J. Meesters, Thibault Debord, Jiří Pešek, Bram Lories, Peter J. Yunker, Hans P. Steenackers, Bart Smeets

## Abstract

Biofilms are structured microbial communities whose extracellular matrix is widely regarded as a basis of their protection against antimicrobial compounds. Yet how matrix production by individual bacteria gives rise to collective architecture and antimicrobial protection remains poorly understood. Here, we systematically varied expression of the master biofilm regulator *csgD* in *Salmonella enterica* and found that increasing matrix production reorganizes biofilms from dense, isotropic packings into sparse, nematically aligned communities by altering cell–cell interactions. By combining experimentally measured biofilm architectures with reaction–diffusion modeling, we show that these structural changes produce distinct patterns of antimicrobial killing, ranging from preferential killing near the liquid-biofilm interface to more uniform killing throughout the community. Consequently, increasing matrix production unexpectedly reduces antimicrobial survival by shifting the biofilm into different transport regimes, while strain-specific physiological differences further modulate antimicrobial depletion. Rather than acting as a passive barrier, EPS therefore shapes antimicrobial susceptibility by reorganizing biofilm architecture and its transport properties. EPS thus provides a physical link between molecular regulation, collective architecture and antimicrobial survival, providing a quantitative framework for understanding how cellular matrix production generates emergent biofilm function.

## Main

Biofilms are multicellular microbial collectives that inhabit and influence virtually every habitat on Earth^1,2^. Their defining hallmark is the production of a matrix consisting of extracellular polymeric substances (EPS) — a composite of proteins, polysaccharides, lipids, and nucleic acids — that underpins many of the emergent functions of biofilms^2^. Among these, protection against antimicrobial compounds is perhaps the most striking and currently problematic, allowing biofilm-associated cells to survive antimicrobial concentrations that are orders of magnitude above those capable of killing their planktonic counterparts^1,3^.

Biofilm-associated protection emerges from the interplay between the intrinsic susceptibility of individual cells and the transport of antimicrobial compounds through the community^4^. In the absence of flow, this transport is governed by the balance between molecular diffusion and local reaction, e.g., consumption, alteration and sequestration by individual cells. Although the EPS matrix is often viewed as the principal barrier to diffusion, diffusivities within biofilms are frequently comparable to those measured in water, suggesting that transport limitations often cannot be understood from matrix permeability alone^5^.

Instead, diffusion and cellular reactions together establish characteristic transport length scales over which chemical compounds are depleted or redistributed. Such transport length scales have been recognized as fundamental determinants of microbial interactions, governing processes including nutrient competition^6–8^, metabolic cross-feeding^9,10^, quorum sensing^11^, and antimicrobial tolerance of microbial communities^12^. Because these transport processes unfold within a spatially organized community, the arrangement of cells and matrix may itself influence the chemical environment experienced by individual cells. Yet we often lack a quantitative understanding of how EPS is organized within biofilms and how its production shapes community architecture, leaving unclear whether and how this architecture feeds back onto antimicrobial transport and collective protection.

Biofilm architecture emerges from local mechanical interactions between constituent cells and their surrounding environment^13,14^. Biophysical approaches have begun to reduce the biochemical complexity of the EPS matrix to fundamental physical interactions, such as adhesion, friction, and steric repulsion. Within this framework, EPS matrix acts as the critical physical mediator, translating molecular regulation into collective organization through localized cell-cell potentials^14–18^. Changes in EPS production could therefore propagate from the molecular scale, through altered cell–cell interactions, to the spatial organization of the community. Yet how these EPS-mediated mechanical interactions translate into differences in antimicrobial survival remains unclear.

In prominent members of the *Enterobacteriaceae* family—including major pathogens such as *Es-cherichia*^19^, *Citrobacter*^20^, and *Salmonella*^21^—the spatial organization of the biofilm is orchestrated by the biofilm master regulator CsgD^22^. This transcription factor coordinates the synthesis of the two primary matrix components: curli amyloid fibers and phosphoethanolamine cellulose, which serve as the structural scaffold for these enteric biofilms^22–24^. Using *Salmonella enterica* as a model, we systematically vary *csgD* expression to modulate matrix abundance. By combining quantitative single-cell imaging, mechanically coarse-grained active-matter simulations, and reaction–diffusion modeling, we resolve how changes in EPS production propagate across spatial scales, from molecular regulation and local cell-matrix interactions to three-dimensional biofilm architecture and antimicrobial survival. We show that matrix production controls antimicrobial survival through a physical cascade: molecular regulation sets the spatial range of intercellular interactions; these interactions sculpt the three-dimensional architecture of the biofilm; and this emergent architecture establishes the reaction-diffusion regime that governs antimicrobial exposure, with metabolic activity further modulating local antimicrobial depletion within these architectural contexts to ultimately determine collective survival.

### *csgD* expression controls individualized EPS envelope thickness

To dissect how genetic alterations in matrix production manifest at the single-cell level, we engineered a matrix-production system in *Salmonella enterica*, by introducing a synthetic library of constitutive Pro-series^25^ promoters in an EPS-deficient Δ*csgD* mutant to drive *csgD* expression. While the native WT promoter exhibits its characteristic bistable expression during the biofilm phase^26^, the engineered Pro-series strains displayed a spectrum of monostable, constitutive expression levels that broadly overlap with the WT range (Fig. 1a). Across all *csgD*-expressing strains, we observed that the two primary EPS components, curli amyloids and pEtN-cellulose, remain highly localized around individual cells, decaying into the extracellular space (Fig. 1b–c). For all bacteria in our imaging volume, we quantified the local EPS density from the volume-integrated fluorescence intensity and the physical EPS envelope thickness from the characteristic decay length of the signal.

**Figure 1:**
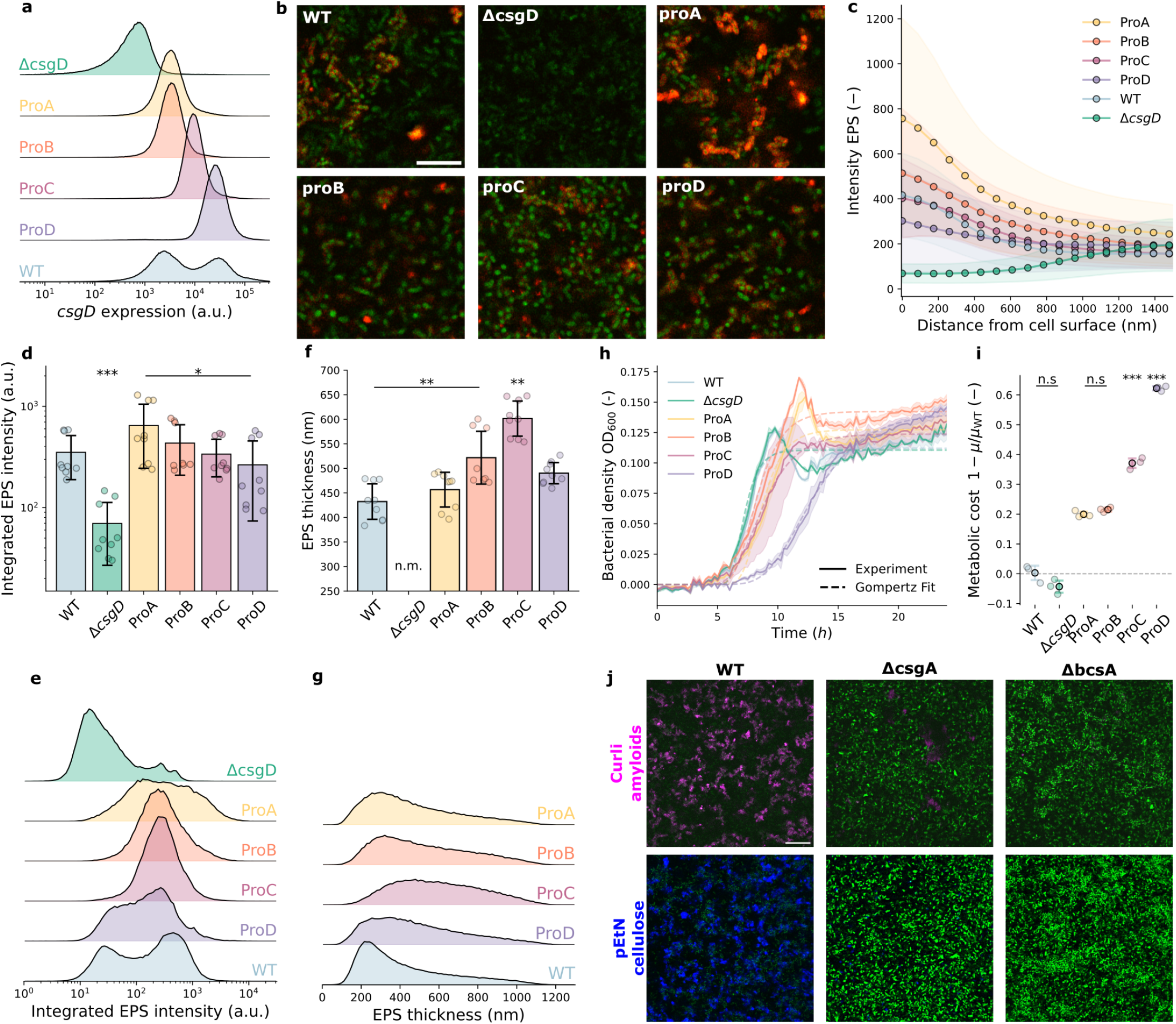
In situ quantification of EPS production as a function of *csgD* expression. In situ quantification of EPS production as a function of *csgD* expression. **a**, We mapped single-cell *csgD* expression levels using a transcriptional sfGFP reporter, demonstrating a spectrum of increasing, monostable expression for the engineered Pro-series and a characteristic bistable pattern for the WT strain. The Δ*csgD* mutant acts as a non-fluorescent control. **b**, Confocal microscopy reveals that the exported EPS matrix (red), stained by EbbaBiolight 680nm, remains tightly bound to individual GFP-expressing cells (green). Contrast settings are identical across all images to allow direct visual comparison. Scale bar, 20 µm. **c**, The EPS optotracer fluorescence intensity decays rapidly away from the bacterial cell surface. Profiles are averaged first over all cells within an individual biofilm; solid lines represent the ensemble average across independent biofilm replicates, and the shaded regions indicate s.d. **d**, The average volume-integrated EPS intensity per cell remains relatively constant across all *csgD*-expressing strains (representing the ensemble average of individual biofilm means). **e**, The distribution of volume-integrated EPS across all biofilm replicates displays a monostable pattern for the engineered Pro-series and a bimodal pattern for the WT, closely mirroring the expression profiles shown in **a. f**, The average thickness of the EPS envelope scales progressively with *csgD* expression (representing the ensemble average of individual biofilm means); due to the lack of localized EPS on the Δ*csgD* mutant, its decay length could not be measured (n.m.). **g**, The corresponding distribution of single-cell envelope thicknesses (length scales) across replicates, shifting progressively toward larger EPS envelopes as *csgD* expression increases. **h**,**i**, Increased *csgD* expression imposes slower growth during planktonic cultivation (**h**) (profiles show means of *n* = 3 independent replicates [solid lines], s.d. [shaded regions], and representative Gompertz fits [dotted lines; see Fig. 1]), resulting in a substantial growth-rate reduction, which we use as a measure of metabolic cost (1 − *µ/µ*_WT_) (**i**). **j**, Targeted staining of curli amyloids (Congo red, pink) and pEtN-cellulose (calcofluor white) in the wild-type and single-knockout mutants (Δ*csgA* and Δ*bcsA*) confirms the physical co-localization and structural synergy of both matrix components. Data show means ± s.d. of *n* = 9 independent biofilms, unless specified otherwise. Statistical significance of pairwise comparisons was determined using a one-way analysis of variance (ANOVA) followed by Tukey’s honestly significant difference (HSD) post-hoc test (^***^*P* < 0.001, ^**^*P* < 0.01, ^*^*P* < 0.05; n.s., not significant).

Although the average EPS density did not differ substantially among the *csgD*-expressing strains (Fig. 1d), the WT population displayed a clear bimodal density distribution that mirrors its underlying genetic bistability (Fig. 1e). In contrast to EPS density, the average physical thickness of the EPS envelope scaled progressively with increasing *csgD* expression (Fig. 1f), with the notable exception of the strongest promoter, ProD. Although the wild-type promoter activity did not yield a clearly bimodal distribution of physical envelope thickness, this structural bistability may remain obscured by the inherently Gamma-like distributions of EPS thickness at the single-cell level (Fig. 1g). We find that increasing *csgD* expression carries a progressive metabolic cost (Fig. 1h-i, SI Fig. 1). We hypothesize that this metabolic stress may become limiting in the hyper-expressing ProD strain, contributing to the reduced envelope thickness relative to the overall trend. To confirm that our measurements accurately captured both key matrix components, we utilized specific dyes to target curli amyloids (Congo red) and pEtN-cellulose (calcofluor white) individually (Fig. 1j). The optotracer signal co-localized with both components in the WT, whereas the curli-deficient (Δ*csgA*) and cellulose-deficient (Δ*bcsA*) single-knockout mutants lacked their respective signals, reinforcing the physical co-localization and structural synergy of both matrix components (Fig. 1j). Together, these single-cell measurements show that increasing *csgD* expression does not primarily increase the density of EPS matrix produced, but instead expands individualized EPS envelopes.

### Individualized EPS envelope thickness drives a collective structural transition

To determine how changes in individualized EPS envelopes control biofilm architecture, we quantified the three-dimensional structure of biofilms formed by strains with different levels of *csgD* expression (Fig. 2a). For each bacterium, we computed the local packing fraction and local nematic order, two geometric descriptors that capture biofilm organization across diverse species^15^ (Fig. 2b, SI Fig. 2). As *csgD* expression increased, the local cellular packing fraction decreased whereas local nematic order increased. This inverse relationship between cellular packing density and cell alignment is the opposite of expectations from soft matter physics, where colloidal particles transition from isotropic to nematic phases with increasing packing fraction^27^. Thus, this result suggests that thicker EPS envelopes, associated with higher *csgD* expression, increase the effective range of cell–cell interactions and the effective cell-plus-EPS packing fraction, thereby strengthening excluded-volume–mediated orientational coupling between neighboring cells. Thus, increasing EPS production unexpectedly generates biofilms with cells that are simultaneously sparser and more aligned, indicating that nanoscale changes in individualized envelopes are amplified into collective structural transitions.

**Figure 2:**
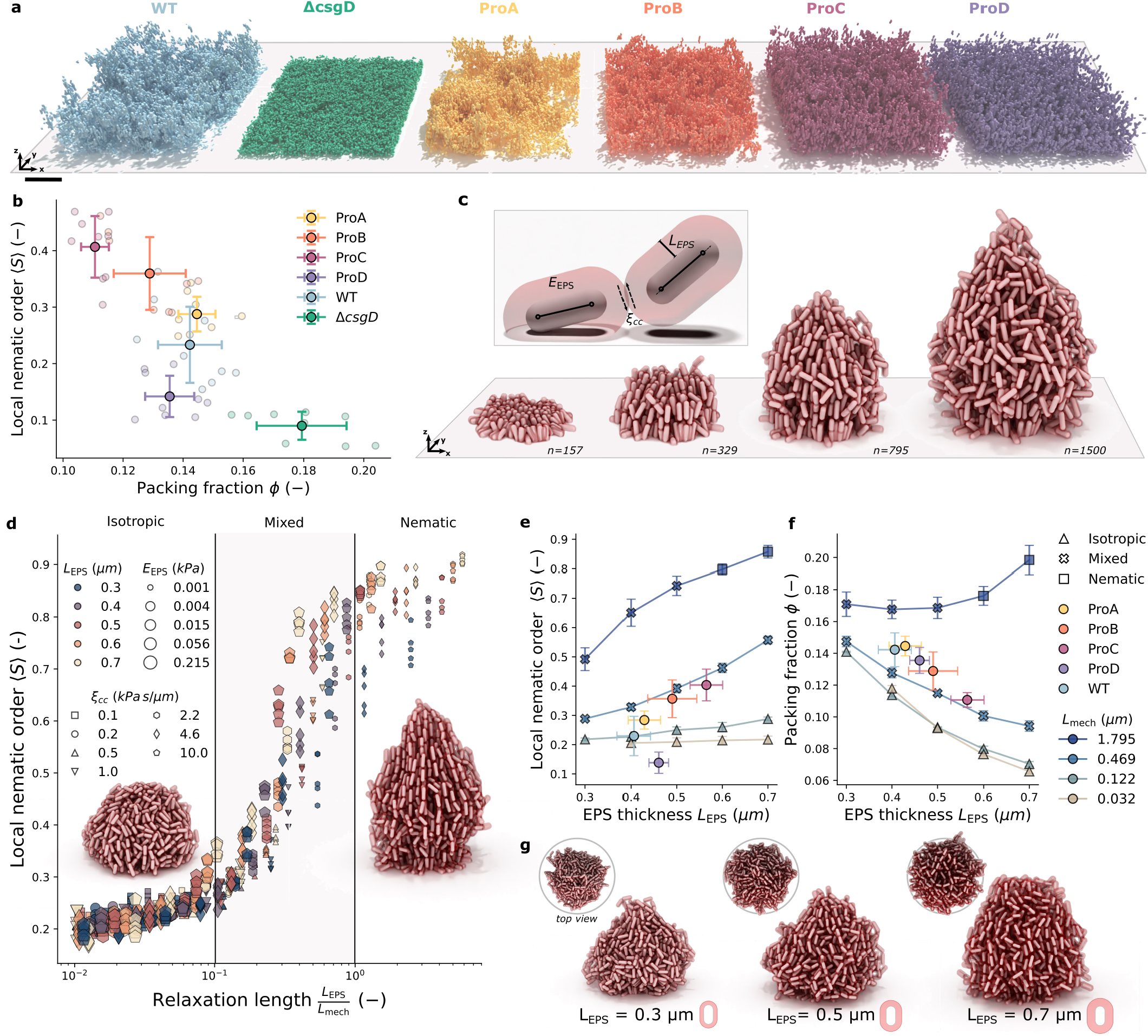
EPS envelope thickness governs the transition from ordered to disordered collective biofilm architecture. **a**, We characterized the architectural changes of experimental biofilms, finding a progressive transition from dense disordered structures to sparser, yet more aligned structures as *csgD* expression increases. **b**, EPS-producing strains systematically diverge from the dense, ordered configurations of non-producing controls as a function of cellular packing fraction (sparseness). **c**, Our individual-based physical model simulates biofilm development over time through cell division, treating bacteria (white) as active particles interacting through their surrounding viscoelastic EPS envelopes (red). **d**, Simulated local nematic order collapses onto a single universal master curve when scaling the physical envelope thickness (*L*_EPS_) by the active hydrodynamic length scale (*L*_mech_), which is parameterized by the EPS stiffness (*E*_EPS_), inter-envelope viscous friction (*γ*_cc_), and the bacterial doubling time (*τ*_g_). see Fig. 3 for additional model visualizations and Fig. 4 for Alexander-de Gennes interaction potentials. **e**,**f**, We mapped experimental local nematic order (**e**) and cellular packing fraction (**f**) onto the simulated parameter space, revealing close quantitative agreement between the model and experimental phase behavior, and highlighting that the empirically observed trends can be explained solely by an increase in EPS thickness *L*_EPS_. **g**, Representative simulated biofilm morphologies highlight the transition from disordered architectures to aligned cluster growth under increasing matrix thickness (*L*_EPS_). In *b, e*, and *f*, experimental data points and error bars represent the ensemble average and standard deviation of *n* = 9 independent biofilm replicates. In **e** and **f** simulation data points and error bars represent the average and standard deviation of *n* = 5 independent replicates.

To understand how the effect of EPS is amplified in a proliferation-driven system, we developed an individual cell–based model in which each bacterium is represented as a deformable spherocylinder surrounded by a spherocylindrical EPS envelope of thickness *L*_EPS_ (Fig. 2c, SI Fig. 3, see Supplementary Information S1). In this model, bacterial proliferation, parameterized by the generation time *τ*_*g*_, is the sole driver of structural rearrangements, and envelope–envelope interactions consist of two generic mechanical contributions: a soft steric repulsion with stiffness *E*_EPS_ and an effective wet friction coefficient *ξ*_*cc*_ that sets dissipation during cell–cell contact. Combined, these parameters yield the active hydrodynamic length scale *L*_mech_ = *E*_EPS_*τ*_*g*_*/ξ*_*cc*_, which measures how far growth-induced mechanical stresses propagate through the community before being dissipated by friction. Comparing this scale with the EPS interaction length *L*_EPS_ defines a dimensionless criterion for the propagation of growth-induced disturbances where local disorder emerges only when *L*_mech_ > *L*_EPS_. This length-scale criterion robustly predicts the structural regimes of the biofilm, from disordered to mixed to fully nematic (Fig. 2d). Moreover, this behavior is robust to the precise choice of interaction potential (SI Fig. 4). Hence, treating EPS as individualized per-cell envelopes that mediate soft steric interactions, rather than as a homogeneous viscous matrix, accounts for the experimentally observed increase in sparsity and nematic order with EPS thickness: in the intermediate regime where *L*_EPS_ ≈ *L*_mech_, increasing *L*_EPS_ shifts the system toward nematic alignment and reduced packing, in quantitative agreement with our measurements (Fig. 2e–g). These results establish biofilm architecture as the emergent physical consequence of individualized EPS production.

**Figure 3:**
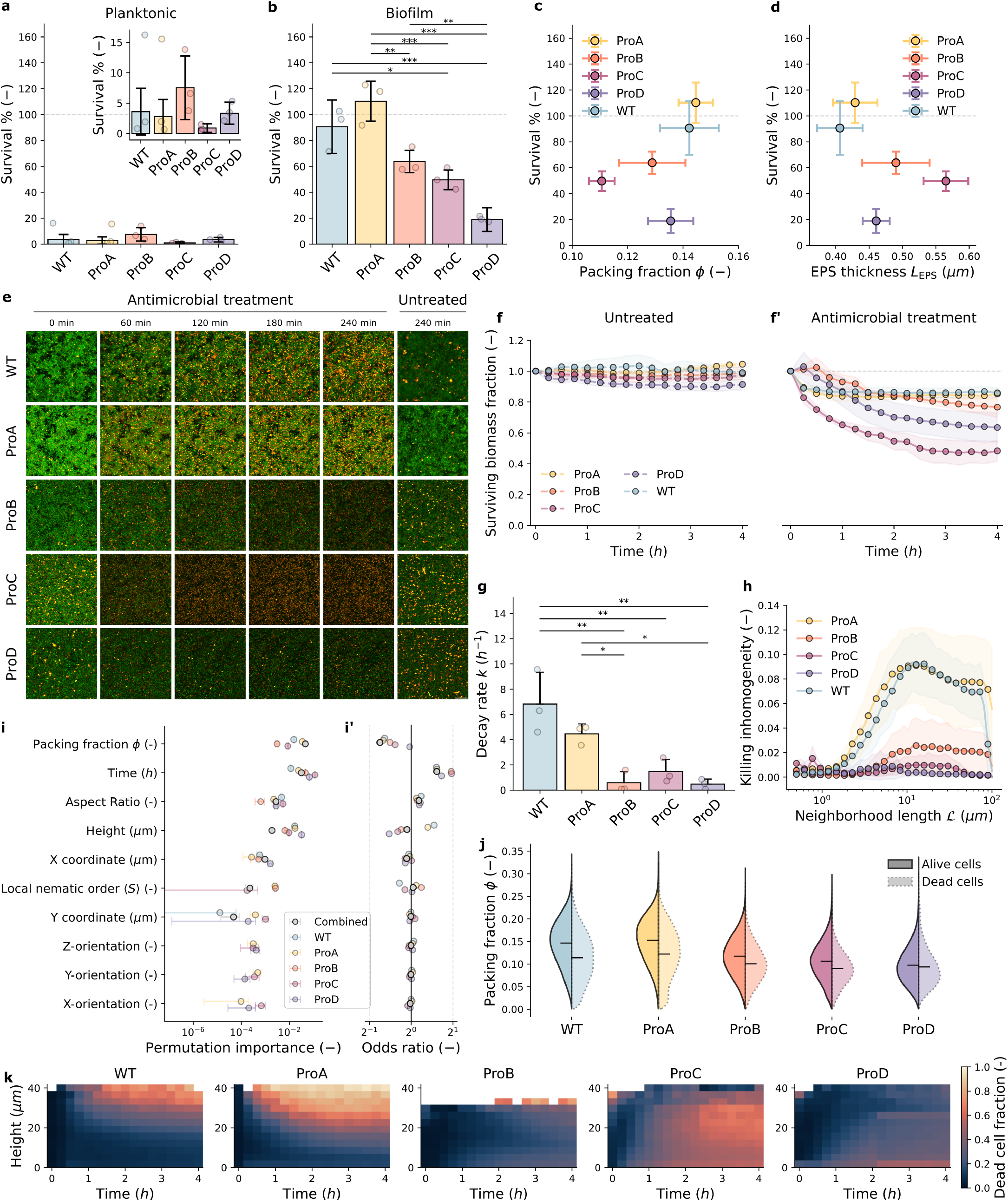
The effect of biofilm architecture on antimicrobial susceptibility. **a**, We observed no differences in survival within the planktonic phase. **b**, The biofilm phase, however, exhibits enhanced survival compared to its planktonic counterpart and shows a decrease in survival with increasing *csgD* expression. **c, d**, Biofilm survival correlates with the average biofilm packing fraction (**c**) rather than with the average thickness of the EPS envelope, *L*_EPS_ (**d**). **e**, Using confocal microscopy, we evaluated the biofilm response in the presence and absence of antimicrobial treatment (500 µg mL^−1^ CTX for 4 h). Representative time-series average projections of biofilms constitutively expressing GFP (green) and stained with propidium iodide (PI, red) reveal the spatial distribution of membrane-compromised cells (see Supplementary Fig. 8). Scale bar indicates 20 *µ*m. **f, f’**, In the absence of treatment, biofilm biomass remains constant (**f**), whereas we observe exponential decay over time in the presence of antimicrobial treatment (**f’**), reconstructed from segmented confocal time-lapses. **g**, This biomass decay rate is high for the WT and ProA strains, whereas we observed significantly slower decay rates for the ProB, ProC, and ProD strains. **h**, Spatial killing heterogeneity varies as a function of the neighborhood length scale *l* under untreated and treated conditions (see Supplementary Fig. 9). **i, i’**, Logistic regression models predict single-cell death across strains, showing standardized feature importance (permutation importance) ranked on the predictive model for all pooled strains (**i**) and the corresponding odds ratios for cell death (**i’**) (see Supplementary Fig. 10). **j**, Surviving and dead cells occupy distinct local packing fraction distributions across the different strains. **k**, Kymographs reveal the spatial killing dynamics as a function of biofilm height over time. Error bars represent s.d. For population-level survival (**a**–**d**), we performed three independent biofilm assays with three technical repeats each (*n* = 9 independent biofilms). Architectural metrics represent the mean ± s.d. of nine independent biofilms (*n* = 9). Microscopic temporal characterization of survival was performed on three independent biofilms (*n* = 3). Statistical significance of pairwise comparisons was determined using a one-way analysis of variance (ANOVA) followed by Tukey’s honestly significant difference (HSD) post-hoc test (^***^*P* < 0.001, ^**^*P* < 0.01, ^*^*P* < 0.05; n.s., not significant). In **a** and **b**, survival was quantified as the fraction of surviving cells after 4 h of 500 µg mL^−1^ CTX treatment relative to their initial density before treatment.

**Figure 4:**
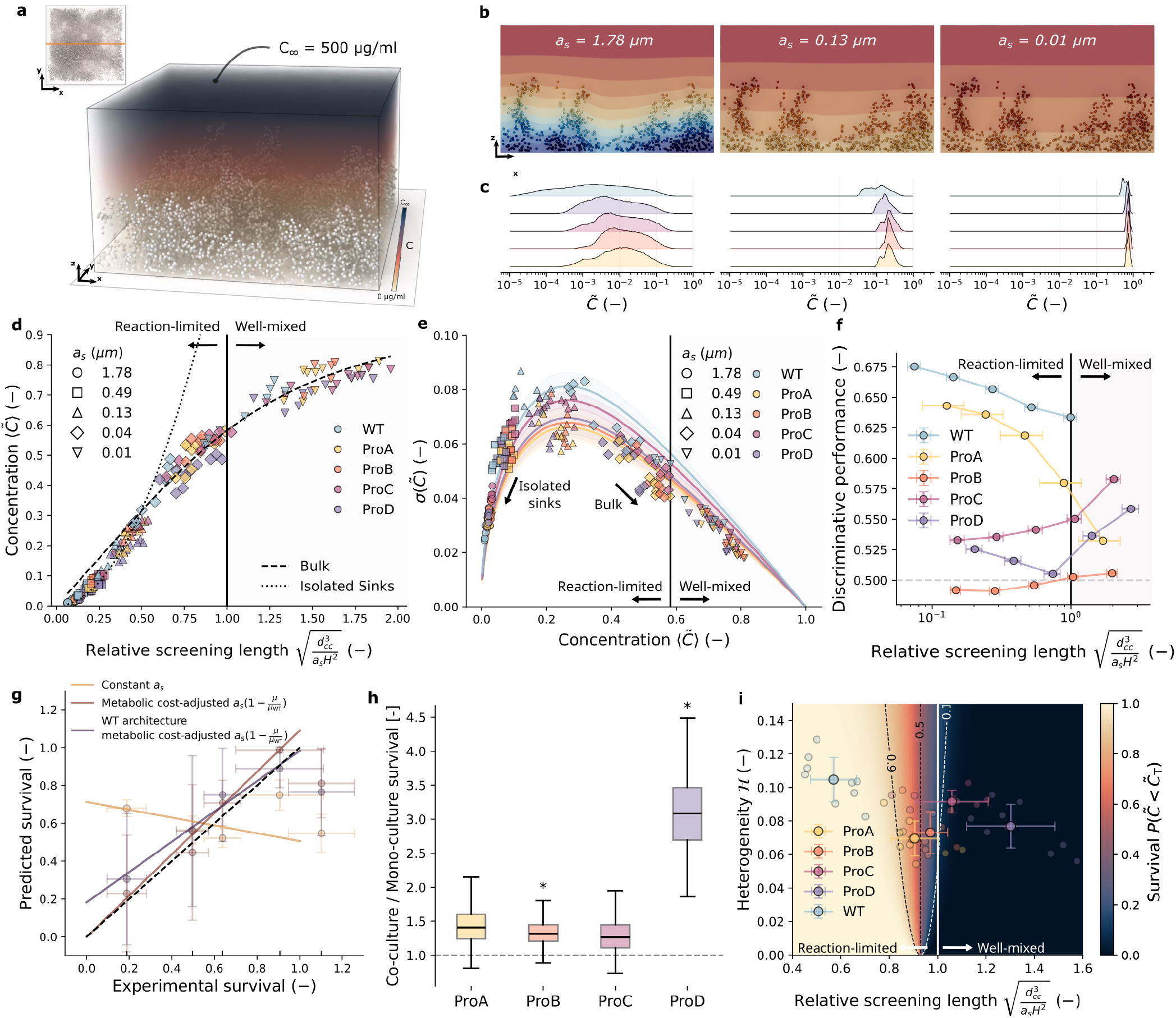
Reaction-diffusion modeling of different biofilm architectures predicts bacterial survival. **a**, We established a reaction-diffusion model to simulate antimicrobial penetration, incorporating experimentally segmented bacterial positions as sinks (white spheres) within the diffusing agent’s concentration field (inset shows the corresponding top-down view of the simulated biofilms and the cross-sectional plane depicted in **b**). **b**, Sliced concentration profiles through wild-type (WT) biofilms show steep local gradients that flatten as the sink strength (*a*_s_ = *k/D*) decreases. **c**, Differences in biofilm architecture significantly alter the normalized concentration distributions 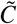 across a wide range of sink strengths (*a*_s_). **d**, We quantified the effective screening length scale based on both architectural properties (average cell-cell distance *d*_cc_ and biofilm height *H*) and sink strength *a*_s_. At low screening lengths (< 0.5), the mean concentration scales as a collection of isolated sinks (dotted line, 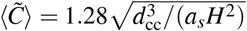, *R*^2^ = 93 within the isolated sink regime), whereas at high screening lengths, it behaves as a continuum bulk (dashed line, 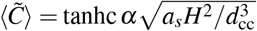, with fitting parameter *α* = 1.58 and *R*^2^ = 0.97 within the bulk regime). Within this bulk regime, we identify a reaction-limited regime where the relative screening length does not span the full biofilm height (< 1) and a well-mixed regime where it does (> 1), demarcated by a solid black vertical line. **e**, To quantify the effects of spatial heterogeneity, we evaluated the standard deviation of concentration 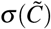 as a function of the mean concentration 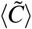. From this relationship, we extracted the architectural heterogeneity *ℋ* using a single-parameter fit (see Supplementary Fig. 15 and Supplementary Information S4). **f**, We evaluated the predictive power of simulated concentration fields on local cell death using logistic regression models. **g**, We evaluated which sink strengths *a*_s_ and killing thresholds best approximate the bulk population survival. We found that architecture, when combined with a metabolic cost-based adjustment of effective sink strength (cf. Fig. 1i), generates the best overall agreement (*R*^2^ = 0.98, slope = 1.09, intercept = 0.00) compared to architecture alone (*R*^2^ = 0.12, slope = −0.21, intercept = 0.71) or pure metabolic adjustment (*R*^2^ = 0.96, slope = 0.80, intercept = 0.18). **h**, We observed that growing strains in co-culture with Δ*csgD* generally increases resilience against antimicrobial treatment compared to the same strains grown in monoculture. **i**, A reaction–diffusion survival landscape maps the best-fitting bulk survival shown in **g**, where the white line indicates the transition from a reaction-limited to a well-mixed bulk regime. Error bars represent s.d. In **b**–**e, g**, and **i**, diffusion simulations were performed on nine independent (*n* = 9) empirical biofilm architectures for each strain. In **f**, we performed logistic regression on three independent microscopy time-lapses (*n* = 3) per strain; discriminative performance was calculated as the mean area under the receiver-operating characteristic curve (AUC-ROC) using a 4-fold cross-validation train/test split bootstrapped 50 times (*n* = 50). In **g**, experimental survival data represent the average ratio of three independent biological repeats (*n* = 3), each consisting of three technical repeats per strain. Statistical significance was determined using a one-sample *t*-test comparing values against a ratio of 1, indicated by the dotted line (^***^*P* < 0.001, ^**^*P* < 0.01, ^*^*P* < 0.05).

### Collective architecture overrides localized EPS protection

The architectural transition raises a central question: does reorganizing the physical structure of the biofilm also change its protective function? We therefore treated both the planktonic and biofilm phases with cefotaxime (CTX) at concentrations far above the WT minimum inhibitory concentration (500 µg mL^−1^ ≈ 4000-fold) (SI Fig. 5). In the planktonic fraction, neither survival nor MIC differed significantly across the *csgD*-expressing strains, indicating that neither altered EPS production nor the metabolic burden of constitutive *csgD* expression affects innate susceptibility to CTX (Fig. 3a and SI Fig. 5, 6). In contrast, in the biofilm phase, survival decreased with increasing *csgD* expression (Fig. 3b), contrary to the common view that increased EPS production improves protection against antimicrobials. Given the constitutive nature of the promoter constructs, the discrepancy between planktonic and biofilm survival suggests that biofilm organization contributes substantially to the antimicrobial response, beyond any localized effects of thicker EPS envelopes (Fig. 3c-d). To investigate biofilm survival dynamics, we tracked biofilm architecture over time (Fig. 3d-f). In the absence of CTX, viable biomass remained largely stable, except in the highest EPS-producing strain, ProD (Fig. 3f). Upon CTX exposure, all strains showed an approximately exponential decline in viable biomass (Fig. 3f’), but with distinct dynamics: ProA and WT displayed rapid initial killing followed by high residual survival, whereas the higher EPS-producing strains exhibited slower decay but lower final survival fractions (Fig. 3g). The endpoint survival measured by microscopy correlated with population-level CFU quantification (SI Fig. 7).

To quantify antimicrobial killing at single-cell resolution, we classified membrane-compromised cells by propidium iodide (PI) staining (SI Fig. 8). To determine how this biofilm-mediated antimicrobial response is spatially organized within the biofilm, we quantified the heterogeneity of the local killing probability conditioned on local cell density as a function of the neighborhood length scale, *l* (see Supplementary Information S3). In untreated conditions, all strains exhibited a distinct maximum at *ℒ* ≈ 5–10 µm, indicating that spontaneous cell death is not spatially random but reflects the underlying biofilm microstructure (SI Fig. 9). Following antimicrobial treatment, the strains displayed markedly different behaviors (Fig. 3h). In WT and ProA biofilms, the density-dependent killing heterogeneity increased substantially, reaching a broad maximum around *l* ≈ 10 µm and remaining high over a wide range of larger length scales. This indicates that biofilm architecture at larger spatial scales remains strongly associated with the local killing rate, consistent with a macrostructural contribution of biofilm organization to antimicrobial susceptibility. In contrast, the remaining strains exhibited a progressively weaker heterogeneity signal with increasing *csgD* expression, indicating that killing becomes more homogeneous and less associated with local density at the scale of the biofilm structure.

To identify which architectural features underlie the observed spatial organization of killing, we related cell survival to local geometric and temporal descriptors using logistic regression (SI Fig. 10). Local biofilm architecture was predictive of cell fate, both across the pooled dataset and within individual strains. Across all strains, local packing fraction emerged as the dominant architectural predictor, together with treatment time: cells in sparser local environments and subjected to longer antimicrobial exposure are more likely to die (Fig. 3i-i’). In contrast, the influence of vertical position differs between strains. WT and ProA cells are predicted more likely to die near the biofilm surface, whereas ProB, ProC, and ProD exhibit the opposite trend (Fig. 3i’). These local relationships are reflected in the distributions of packing fractions occupied by surviving and dead cells (Fig. 3j, SI Fig. 11). Consistent with these differences, kymographs of cell survival reveal distinct spatial killing patterns across the biofilm, with WT and ProA exhibiting predominantly top surface-localized killing, whereas ProB, ProC and ProD display a more homogeneous distribution of cell death throughout the biofilm depth (Fig. 3k, SI Fig. 12, 13). These observations suggest that biofilm architecture may influence antimicrobial susceptibility by shaping the local environment experienced by individual cells.

### Position and metabolic activity jointly govern antimicrobial survival in structured biofilms

The dependence of survival on local biofilm architecture prompted us to test whether differences in antimicrobial exposure could account for the observed killing patterns. We therefore simulated steady-state antimicrobial transport in the measured biofilm architectures (Fig. 4a, SI Fig. 14). Cells are represented as reactive point sinks with sink strength *a*_*s*_ = *k/D*, where diffusion (*D*) competes with sequestration (*k*), determining the spatial extent of local depletion (Fig. 4b–c). To classify biofilm architectures according to their governing reaction-diffusion regime, we solved the steady-state diffusion equation over a range of sink strengths at the measured cell positions. The resulting cell-averaged concentration, 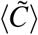, collapses onto a single curve when plotted against the relative screening length, 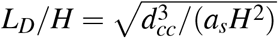, where *d*_*cc*_ is the mean cell–cell spacing and *H* is the biofilm height (see Supplementary Information 17). Small screening lengths yield an isolated-sink regime governed by local depletion around individual cells. As the screening length increases, the system approaches a continuum bulk regime, which can be further divided into a reaction-limited regime, where the screening length remains shorter than the biofilm height, and a well-mixed regime, where it exceeds the biofilm height (Fig. 4d)

Predicting transport-mediated differences in survival further requires quantifying the heterogeneity of exposure across cells. Across the measured biofilm architectures, the cell-to-cell variability in simulated concentration follows a bulk approximation with the mean concentration (Fig. 4e; Supplementary Information S4). The mean concentration determines the characteristic scaling of variability, while biofilm architecture modulates its magnitude through a single structural heterogeneity parameter (SI Fig. 15). Consequently, variability is minimal in both the isolated-sink and well-mixed limits, where cells experience similarly depleted or homogeneous environments, and maximal in an intermediate regime where depletion zones partially overlap. Incorporating this heterogeneity into a survival model reveals a survival advantage for structurally disordered biofilms at low killing thresholds, as spatial fluctuations create protected environments even at fixed mean concentrations (SI Fig. 15).

To test whether these inferred reaction–diffusion regimes explain the observed spatial killing patterns, we fitted logistic models relating the simulated antimicrobial concentration at individual cell positions to the experimental survival over a range of relative screening lengths (Fig. 4f, SI Fig. 16). Predictive performance was highest for WT and ProA at screening lengths below one, consistent with a reaction-limited regime, indicating that local antimicrobial depletion is predictive of cell fate in these architectures. In contrast, ProB, ProC, and ProD showed only weak predictive performance across all screening lengths, with the highest performance occurring in the well-mixed regime, suggesting that structural effects contribute little to the observed killing patterns. However, differences in biofilm architecture alone do not reproduce the observed bulk survival differences across strains (Fig. 4h, SI Fig. 17), indicating that the effective transport parameter *a*_*s*_ must also vary between conditions. We next asked whether incorporating strain-specific differences in effective antimicrobial uptake could account for the remaining variation in survival. We therefore use planktonic growth rate not as a direct measurement of antimicrobial uptake, but as a physiological proxy with which to rescale the effective sink strength *a*_*s*_ between strains. This single-parameter model better predicts bulk survival compared with a constant *a*_*s*_ across all strains (Fig. 4g, SI Fig. 18– 19), consistent with growth-associated metabolic activity providing a useful proxy for strain-specific differences in effective antimicrobial uptake. To further test whether differences in effective sink strength are associated with metabolic activity, we co-cultured the fast-growing, EPS-deficient Δ*csgD* strain with the respective strains, thereby perturbing the metabolic environment while aiming to preserve biofilm architecture (Fig. 1i). Relative to monoculture, co-culture increased survival across strains, with the strongest effect observed for ProB and ProD (Fig. 4h). These results are consistent with metabolically more active cells contributing to local antimicrobial depletion and thereby altering the effective transport environment experienced by neighboring cells.

Using the screening lengths inferred from the survival analysis (Fig. 4f-g) together with the structural heterogeneity from the bulk approximation (Fig. 4e), we position each strain within a two-dimensional reaction–diffusion survival landscape (Fig. 4i). WT and ProA fall within the reaction-limited regime, where local depletion generates protected microenvironments, whereas ProB, ProC and ProD are in the well-mixed regime, where the concentration field is more homogeneous. Hence, although metabolic activity does not measurably alter antimicrobial susceptibility in the planktonic phase (Fig. 3a), its effects become consequential within biofilms whose architecture places them in distinct reaction-diffusion regimes. The predominantly vertical survival contours indicate that the reaction-diffusion regime dominates survival, while architectural heterogeneity provides a secondary structural contribution (Fig. 4i).

## Discussion

Biofilm matrix production is widely regarded as a defining determinant of biofilm function, yet how matrix production by individual bacteria gives rise to the notorious biofilm-associated antimicrobial protection has remained unclear^4^. Under the prevailing paradigm, the matrix is conceptualized as a physical shield that slows down antimicrobial penetration, implying that increased matrix production inherently yields a more effective barrier. However, it has long been argued that treating the matrix as a simple diffusion barrier fails to account for the magnitude of biofilm tolerance^5,28^. Here, we show that antimicrobial susceptibility is governed not simply by the amount of extracellular matrix produced, but by the physical architecture that matrix production generates. Progressive thickening of individualized amyloid–cellulose envelopes drives a transition from densely packed isotropic communities to sparser nematically aligned biofilms, reorganizing the chemical environment experienced by individual cells. Thus, rather than acting as a passive diffusion barrier, the biofilm matrix determines antimicrobial survival indirectly by shaping the collective physical organization of the community.

Our experiments and simulations converge on a biophysical picture in which the biofilm EPS primarily modifies intercellular interactions, rather than acting as a continuous background medium. This challenges the common view of the matrix as a continuous adhesive slime surrounding the community. Correspondingly, biofilm architecture emerges from the balance between growth-driven mechanical stresses and the localized interactions imposed by EPS envelopes^14,15,17,18^. More broadly, this places EPS-mediated local interactions within a physical framework in which changes in these interactions can drive transitions in biofilm architecture, including transitions between attached and dispersed states, and potentially shape interactions between competing populations^29–32^. Although the adopted physical description is agnostic of the detailed molecular identity of the matrix, our experiments focus on the curli–cellulose matrix of *S. enterica*. Whether other EPS chemistries generate comparable interaction ranges and architectural transitions remains to be tested. Importantly, we show that architectural changes can become functionally relevant by determining the transport regime in which the community operates. In this view, biofilm architecture is not simply a structural phenotype but a physical determinant of chemical transport setting the balance between transport and consumption. Consequently, the influence of emergent architecture on antimicrobial exposure is itself regime dependent, becoming strongest when transport and uptake occur over comparable length scales^33^. Because this balance depends only on the relative scales of molecular transport and community organization, the framework should extend beyond antimicrobials to nutrients, metabolites, signaling molecules and other diffusible compounds exchanged within microbial communities.

A complementary consequence is that antimicrobial susceptibility cannot be understood by considering bacterial physiology independently of community organization. Reduced uptake is commonly regarded as a route to individual survival because it limits intracellular drug accumulation^34^. Our results instead suggest that differences in metabolic activity can further influence collective antimicrobial exposure, although we note that rescaling sink strength using growth rate is phenomenological rather than a direct measurement of antimicrobial uptake. Our reaction-diffusion framework intentionally coarse-grains these processes into an effective sink strength and therefore does not distinguish between intracellular uptake, degradation, sequestration, or other mechanisms of antimicrobial removal. This abstraction allows us to isolate how spatial organization and effective depletion interact, while identifying the molecular processes underlying strain-specific sink strengths will require direct measurements of antimicrobial uptake and transformation. While the ‘reaction-sink’ concept is classically associated with enzymatic degradation, such as *β*-lactamases^35,36^, our findings are consistent with metabolic activity contributing to antimicrobial depletion, analogous to collective transport bottlenecks observed in larger systems^33^. Whether greater cellular antimicrobial uptake sensitizes or collective antimicrobial removal protects a population therefore depends on spatial context. Thus, metabolic activity may have opposite consequences at the individual and community levels, with increased cellular demand increasing susceptibility through greater intracellular exposure while collective consumption may reduce exposure for neighboring cells. Our experiments probe these effects at a defined CTX concentration, and changing the antimicrobial concentration or other treatment parameters (e.g. antimicrobial concentration, identity, or exposure conditions) may alter the relative contributions of collective architecture and single-cell responses to killing. Nevertheless, the underlying length-scale formulation is not specific to these conditions and provides a basis for testing how these parameters reshape the survival landscape.

Our findings redefine the role of the biofilm matrix. Rather than functioning primarily as a passive protective barrier, EPS production directly alters biofilm architecture, thus organizing microbial communities into distinct physical states that determine how chemical gradients emerge and how antimicrobial exposure is distributed among individual cells. More generally, our results establish a hierarchy of coupled physical length scales linking gene regulation to community function, in which genetic changes in EPS production alter single-cell interaction ranges, collectively reshape biofilm architecture and its reaction-diffusion transport regime, and ultimately influence antimicrobial survival. More broadly, this provides a physical framework for understanding how local cellular properties propagate across scales to generate emergent functions at the level of microbial communities.

## Author contributions

B.S., H.S. and T.B. conceptualized the project. B.L., H.S., J.M., T.B. and T.D. designed the experiments. J.M. and T.D. performed the experiments. B.L., J.M. and T.B. analyzed the experiments. B.S., J.P. and T.B. developed and performed the simulations. B.S., J.P. and T.B. analyzed the simulations. B.S. and T.B. wrote the first draft of the paper, and all authors contributed to the revision of the manuscript.

## Acknowledgments

TERB was funded by a grant from the Research Foundation – Flanders (FWO) (grant no. 1266425N and V414424N). TD was funded by a grant from Research Foundation – Flanders (FWO) (grant no. 11PV924N). PJY was funded by a grant from the National Institutes of Health, National Institute of General Medical Sciences (grant no. 1R35GM138354-01). BS and HPS were funded by a grant from the Research Foundation – Flanders (FWO) (grant no. G043726N) and a grant from the KU Leuven (grant no. C14/22/077). We thank R. Belpaire for comments on the logistic regression; O. Steele-Mortimer, S. Copley, D. Mavridou and B. Van den Bergh for contributing plasmids.

## Data availability

The data underlying all figures is publicly available at Zenodo (doi: https://doi.org/10.5281/zenodo.22940408). All scripts used for figure generation are publicly available at GitLab repository. Raw microscopy images are available from the authors upon reasonable request.

## Code availability

The source code for the simulations presented in this study is publicly available through the GitLab repository. For reproducibility, the complete computational environment, including all required dependencies and the solver backend, is provided as a publicly accessible Docker container for the Mpacts software, together with a Singularity image. The Mpacts backend is freely accessible under a non-open-source license and is provided without warranty. Solver documentation and usage instructions are available through the associated GitLab repository.

## Ethics declarations

The authors declare no competing interests.

## Materials and Methods

### 1 Planktonic growth conditions

All *Salmonella enterica* Typhimurium ATCC14028 strains were grown overnight in lysogeny broth (LB, 10 g L^−1^ tryptone, 10 g L^−1^ NaCl, 5 g L^−1^ yeast extract) supplemented with 40 µg mL^−1^ ampicillin at 37 °C with shaking at 200 rpm.

### 2 Biofilm growth conditions

The biofilm assays were performed by inoculating an initial cell density of 2 × 10^7^ cells mL^−1^ in tryptic soy broth (TSB) diluted 20 times and supplemented with 10 µg mL^−1^ ampicillin. Subsequently, biofilms were grown for 24 h at 25 °C sealed with a breathable membrane. For co-cultured biofilms, the total inoculation density was maintained at 2 × 10^7^ cells mL^−1^ with the two strains inoculated at a 1:1 ratio.

For *csgD* expression and colony-forming units (CFU) quantification, biofilms were inoculated in 6-well plates. After the 24-hour incubation period, the planktonic phase was removed by pipette aspiration, and the biofilm phase was scraped off the surface of the well. Both the planktonic and the biofilm phases were transferred to phosphate-buffered saline (PBS, 1.24 g L^−1^ K_2_HPO_4_, 0.39 g L^−1^ KH_2_PO_4_, 8.8 g L^−1^ NaCl) and passed through a 25G-needle five times to disaggregate potential bacterial clusters. Subsequently, the *csgD* expression of the samples was measured using flow cytometry or the CFU was determined by plating out on LB agar overnight at 37°C.

### 3 Mutant construction

All used *Salmonella* strains and primers are shown in Table 1 and 2, respectively. Genomically fluorescently labeled strains were created using a scarless genome editing protocol. A kanamycin (Km) cassette was amplified from pT2SK for insertion in between STM1666 and STM1667. The promoter and fluorescent protein gene, *sfgfp* or *ecfp*, were amplified from the pUltra library. The Km cassette was inserted into the genome via electroporation into *S*. ATCC14028 wild-type (WT) containing the pSLTS plasmid. Electroporation was repeated using the DNA fragment containing the fluorescent gene with recovery on agar supplemented with 100 ng mL^−1^ anhydrotetracycline. Colonies were assessed for re-obtaining Km sensitivity. Correct insertion was determined via PCR and sequencing (Eurofins).

**Table 1:**
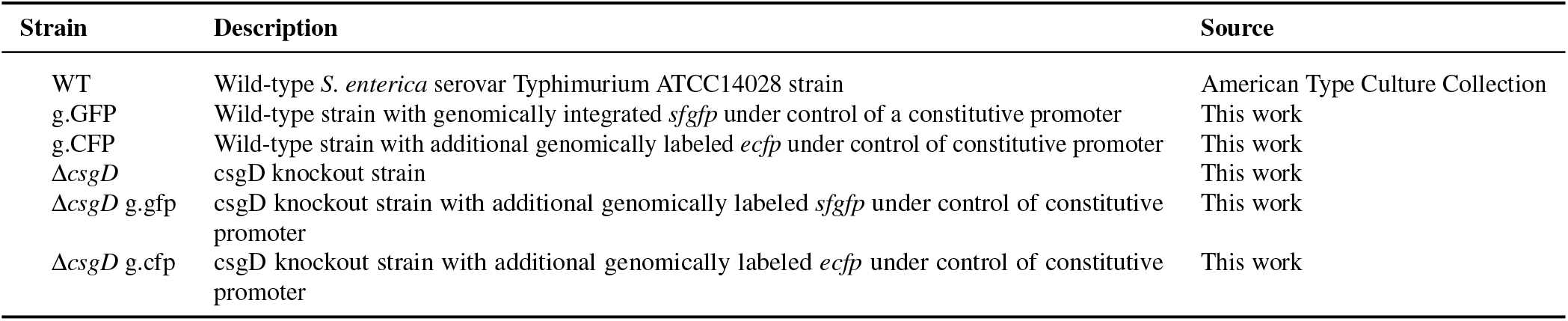
List of strains.

**Table 2:**
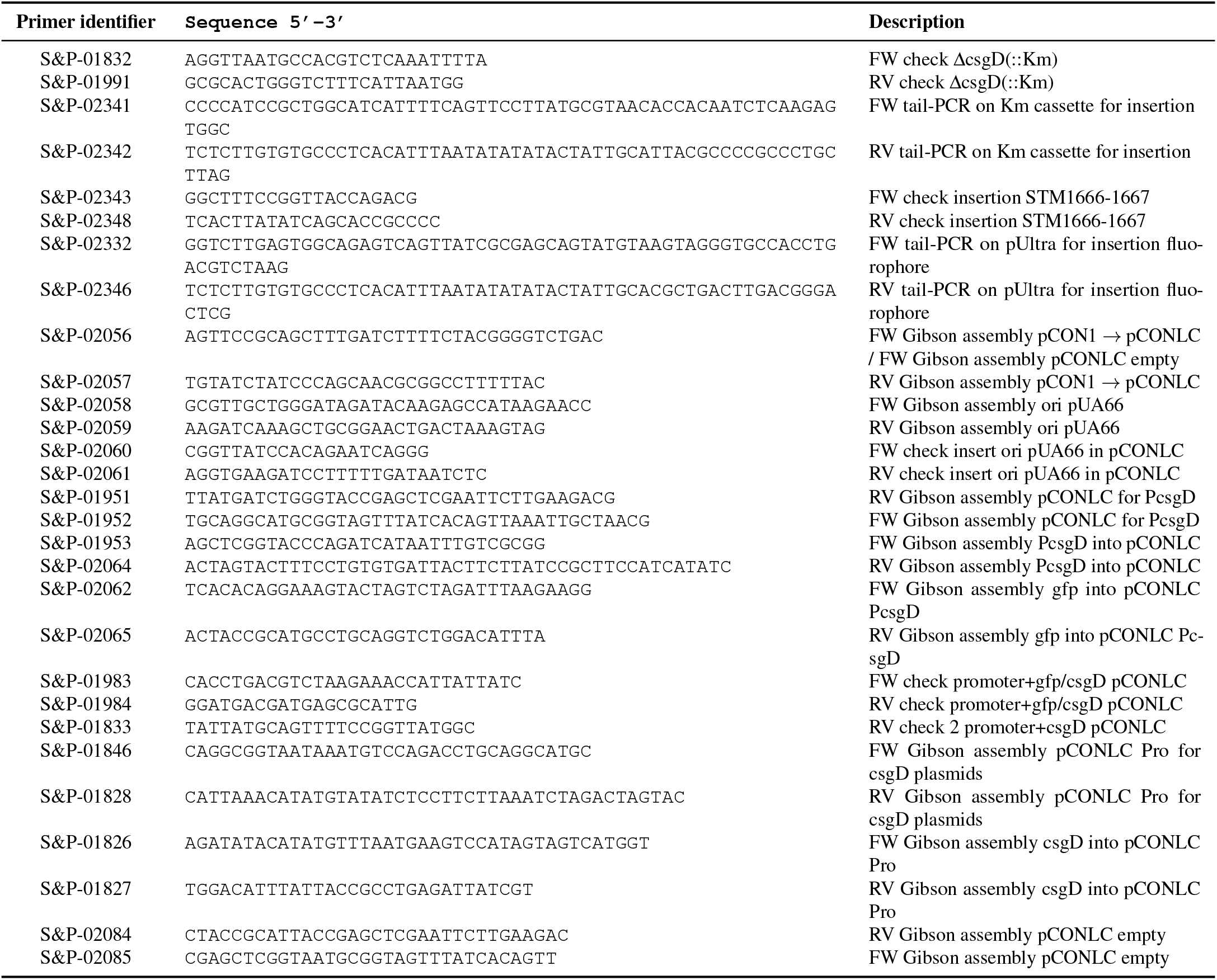
List of primers.

The Δ*csgD* strain was created using the Datsenko and Wanner protocol^37^ with the Δ*csgD*::Km *Salmonella* strain from the McClelland library^38^ as donor for phage transduction. *S*. ATCC14028 WT, g.GFP and g.CFP were used as phage acceptors. Removal of the Km cassette occurred via the introduction of pCP20. Deletion was confirmed with re-obtained sensitivity and sequencing.

### 4 Plasmid construction

An overview of all used plasmids is given in Table 3. The origin of replication (ori) of the pCON1 plasmids was replaced with the ori of pUA66 to obtain the low copy pCONLC plasmid. The *csgD* promoter and gene were amplified from the genome of *S*. Typhimurium ATCC14028. New plasmids were assembled using Gibson assembly. All used primers are shown in Table 2. *E. coli* Top10F’ was used for all cloning steps. Lower antimicrobial concentrations (Ap_10_) were needed for the recovery of pCONLC plasmids.

**Table 3:**
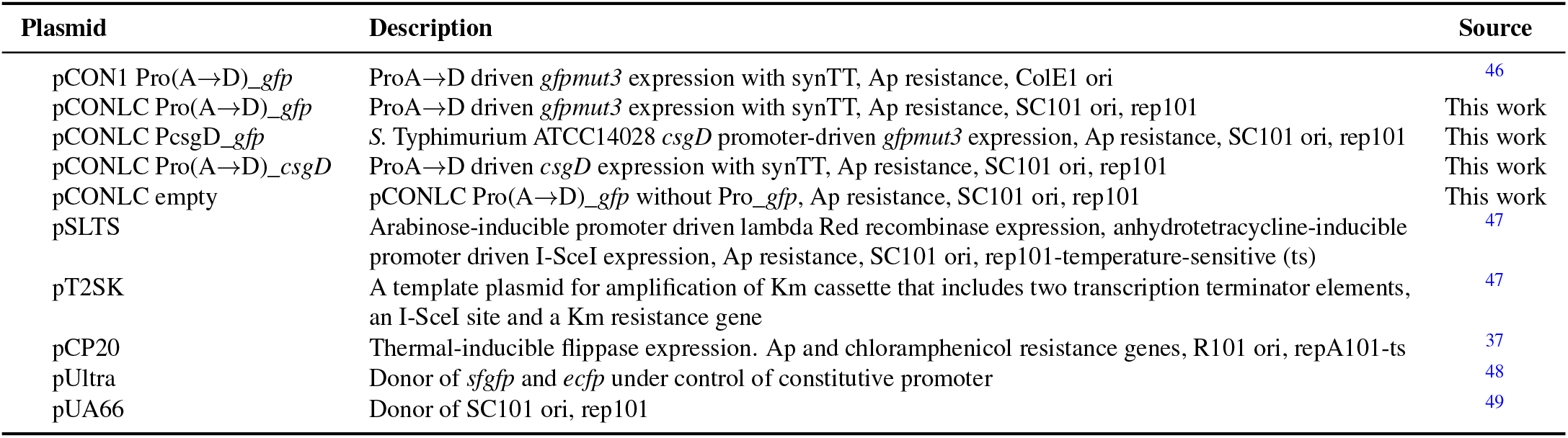
List of plasmids.

### *5 csgD* expression

Expression of a promoter was determined using *gfp* as reporter gene for promoter activity. Samples were measured using a CytoFLEX flow cytometer (Beckman Coulter). Bacteria were distinguished from the background by gating the forward-scatter area (FSC-A) and side-scatter area (SSC-A). Subsequently, doublets were removed from the cell population based on the area (FSC-A) and height (FSC-H) of the FSC signal. Finally, bacterial GFP expression was measured as proxy for *csgD* transcription by excitation with a 488 nm laser.

### 6 MIC determination

Overnight cultures (ONCs) were normalized in 20-fold diluted TSB to a cell density of 4 × 10^7^ cells mL^−1^. Of these normalized cultures, 100 µL was added to each well of a 2-fold serial dilution of cefotaxime in 20-fold diluted TSB in a 96-well plate. The plate was sealed with a breathable membrane and incubated at 25 °C and 200 rpm for 24 h. After incubation, the optical density (OD) at 600 nm of each well was measured using a Synergy MX multimode reader (Biotek). The MIC value was determined per bacterial strain as the lowest antimicrobial concentration where there was no measurable increase in OD as compared to a sterile control.

### 7 Metabolic cost estimation

The ONCs were normalized in 20-fold diluted TSB to a cell density of 4 × 10^7^ cells mL^−1^ in 200 µL volumes of 96-well plate. These cultures were grown for 24 h in Synergy MX multimode reader (Biotek), measuring the optical density at 600 nm every 5 min. The resulting OD curves OD(*t*) were fit per well using the Gompertz model using the Zwietering reparameterization^39^

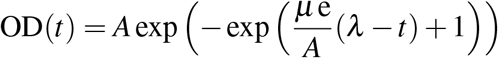

to extract the yield *A*, lag time *λ* and maximum specific growth rate *µ* of each strain. From the maximum specific growth rate the metabolic cost was determined by comparison with the growth rate of the WT strain as 1 − *µ/µ*_WT_.

### 8 Biofilm treatment

After 24 h of incubation, biofilms were treated by adding cefotaxime in TSB to a final concentration of 500 µg mL^−1^ cefotaxime and 1.5 g L^−1^ TSB. For the non-antimicrobial controls, cefotaxime was replaced by demineralized water. Biofilms were treated for 4 h at 25 °C and subsequently plated as described in section 2. The cell count for the treated samples was divided by the cell count at 24 h within the same biological repeat to calculate the antimicrobial susceptibility.

### 9 Image acquisition

Biofilms were imaged in three dimensions using an LSM 880 with an Airyscan detector (Zeiss) using Fast Airyscan mode. The bacterial and EPS signals were obtained by excitation with 488 nm laser and 561 nm, respectively. Images were captured using a 100 × oil immersion objective (alpha Plan-Apochromat 100 × /1.46 oil DIC M27, Zeiss) resulting in a final image size of 84.04 µm by 84.04 µm and voxel dimension 81.7 nm. The Z height of the images was dependent on the height of the imaged biofilm region. The voxel depth was kept constant at 180 nm.

### 10 EPS staining

Biofilms were inoculated in 96-well plates and incubated for 23 h at 25 °C. Subsequently, the biofilm matrix was stained by EbbaBiolight 680 (Ebba Biotech AB), an optotracer previously reported to stain both cellulose and curli^40^, using a final dilution of 1,000-fold. The biofilm was further incubated in the presence of the EPS stain at 25 °C for the duration of 1 h. To localize specific matrix compounds, the biofilms were stained with 50 µg mL^−1^ Calcofluor White (CFW, Sigma-Aldrich) and 40 µg mL^−1^ Congo Red (CR, Sigma-Aldrich) for the detection of cellulose and amyloid, respectively. After incubation for 15 min in the dark, the samples were imaged using the Zeiss LSM 880. CFW-stained biofilms were imaged using the 405 nm laser at a lateral resolution of 100 nm in the x- and y-directions, whereas CR-stained biofilms were imaged using the 561 nm laser at a lateral resolution of 122 nm in both directions.

### 11 Live/dead staining

To evaluate the bacterial killing *in situ*, biofilms were grown as described above and subsequently stained by adding propidium iodide (PI) to a final concentration of 1 µM (Thermo Fisher Scientific). After incubation for 15 min in the dark, biofilms were treated by adding cefotaxime in undiluted TSB to a final concentration of 500 µg mL^−1^ cefotaxime and 1.5 g L^−1^ TSB. For the untreated controls, cefotaxime was replaced by demineralized water. Subsequently, the stained biofilms were imaged for 4 h using FastAiryscan mode, using 488 nm and 561 nm lasers, for the constitutive GFP-expressing cells and PI respectively.

### 12 Image analysis

Images were processed using the Airyscan post-processing (Zen Blue, Zeiss) with automatically determined Wiener filter strength. The confocal Z stacks were scaled to obtain isotropic voxel dimensions of 81.7 nm by 81.7 nm by 81.7 nm. Subsequently, the intensity of each z-slice was normalized by its maximum intensity. The normalized images were filtered by using a three-dimensional Gaussian kernel followed by the Frangi-vesselness filter^41^ (*α* = 0.8, *β* = 0.5, *γ* = 0.01) to select tubular structures. Using Otsu thresholding^42^ on the Frangi-filtered images, a binary mask of the tubular structures was obtained. These binary masks were segmented into separate objects using a watershed algorithm seeded from local maxima in the intensity signal. Foreground and background objects were separated by using Otsu’s threshold on the distribution of mean intensities of all segmented objects. For each object, the length and radius were extracted from the eigenvalues of the inertial tensor of the object. The orientation of each object is obtained from the eigenvector associated with the largest eigenvalue.

#### 12.1 EPS quantification

To quantify the EPS fluorescence surrounding individual bacteria, the segmented bacterial objects were dilated in a voxel-wise manner ranging from the bacterial surface to a maximal length of *L*_max_ = 1.62 µm. The shell-averaged intensity *i*(*x*) of the matrix capsule was determined by integrating the signal over the whole domain normalized with the volume of the shell. The EPS thickness *L*_EPS_ was determined by dividing the first-order moment by the zeroth-order moment 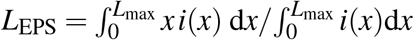, where *x* is the distance from the cell surface.

#### 12.2 Viable/dead classification

Cell viability was assessed for each segmented bacterial object by calculating the ratio between the mean fluorescence intensity of the propidium iodide (PI) channel and the constitutive GFP channel. Cell populations were classified as either PI-positive (dead cells) or PI-negative (viable cells) using a two-dimensional gating strategy based on the PI/GFP intensity ratio and absolute GFP fluorescence.

### 13 Architectural quantification

#### 13.1 Local nematic order

To evaluate the alignment of individual bacteria with their local neighborhood, the nematic order was determined based on the unit direction of the focal bacteria **n**_*i*_ as

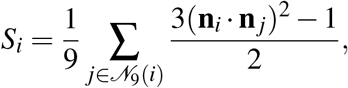

where **n**_**j**_ are the unit directions of the nine nearest neighbors *N*_9_ of the focal bacteria *i*.

#### 13.2 Packing fraction

To determine the local packing fraction *φ*_*i*_, the neighborhood surrounding the focal bacteria was defined as a sphere with radius *R*_*l*_ =5 µm. The local packing fraction was determined by the summation of the convex volumes *V*_*j*_ of bacterial objects, including the focal bacterium, with their centroid within the spherical neighborhood. The packing fraction is then given by normalization by the volume of the neighborhood 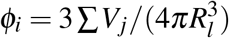. For the matrix packing *φ*_*M,i*_, the volume of the matrix capsule, as determined by the matrix quantification of the images, was considered in addition to the bacterial volumes.

### 14 Quantification of spatial killing heterogeneity

To quantify the dependence of antimicrobial killing on biofilm micro-architecture, we calculated the variance of the mean killing probability across local cell densities. To account for the dependence between spatial resolution and local density in discrete bacterial populations, we evaluated this heterogeneity as a function of the spatial scale *l*. The biofilm was divided into uniformly sampled cubic neighborhoods of size *l*, with local density represented by the number of cells *N* within each neighborhoods. For each *N*, the mean killing probability was calculated across all neighborhoods containing *N* cells. Spatial heterogeneity in killing was then quantified as the variance of these conditional mean killing probabilities as

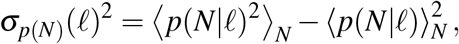

*With* 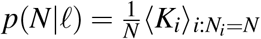, The complete derivation, including the bounds of the variance and its limiting behavior with spatial scale, is provided in Supplementary Note S3.

### 15 Logistic regression

To quantify the predictive power of the measured architectural features, we trained a binary logistic regression classifier using the scikit-learn machine learning library using 4-fold cross validation. On the training portion of the dataset, all continuous predictor variables were standardized and the classifier was optimized using 10^4^ iterations with L2 regularization. To account for class imbalance, class weights were set to the inverse class frequencies. Discriminative performance was assessed using the area under the receiver operating characteristic curve (ROC-AUC), computed from the predicted class probabilities. To quantify the contribution of individual predictor variables, model coefficients were extracted after standardization. Feature importance was additionally assessed using permutation importance. Predictor values were independently permuted 30 times on the held-out test set while keeping the trained model fixed.

### 16 Individual-cell model

Bacteria were represented as deformable spherocylinders defined by two nodes of constant radius connected with an internal spring. EPS envelopes were represented as additional concentric spherocylinders with a larger radius around each cell. Bacterial growth was modeled by linear elongation of the internal spring’s resting length, followed by division whenever the maximum cell length was reached. Interaction between the EPS envelopes of neighbouring cells was represented using an elastic repulsion modeled by the Hertz contact model, as well as viscous friction. The resulting overdamped force balance included spring, viscous friction, repulsive contact and medium-drag forces. Node velocities were obtained by solving the resulting linear system using a conjugate-gradient method, and particle positions were updated using semi-implicit Euler integration. The complete model formulation is provided in Supplementary Note S1 and model parameters in Table 4.

**Table 4:** Simulation parameters for the individual-cell and diffusion model.

| Parameter | Symbol | Value | Units | Rationale/source |
| --- | --- | --- | --- | --- |
| *Cell geometry & growth* |  |  |  |  |
| Cell radius | $R$ | 0.5 | $\mu\text{m}$ | Measured from the average radius of all strains ($0.51 \pm 0.10 \mu\text{m}$). Within the reported range of *Salmonella* radii<sup>50</sup>. |
| Maximum length | $L_{\text{max}}$ | 3.0 | $\mu\text{m}$ | Estimated from empirical length distributions with average bacterial length of ($1.99 \pm 0.70 \mu\text{m}$). |
| Std. dev. of max length | $\sigma_{L_{\text{max}}}$ | 0.15 | $\mu\text{m}$ | Estimated from empirical length distributions. |
| Generation time | $\tau_g$ | 20 | min | Estimated from the planktonic growth rate experiments from the WT strain (Supplementary Fig. 1b). |
| Initial cell density | $\rho_0$ | 0.5 | $\mu\text{m}^{-2}$ | Estimated at the starting point of verticalization<sup>18</sup>. |
| *Mechanical properties* |  |  |  |  |
| Young's modulus of the bacterial cell | $E$ | 1.0 | kPa | Lowered from the reported MPa range<sup>51</sup> for computational stability. |
| Poisson's ratio of the bacterial cell | $\nu$ | 0.45 | – | Within the typical range for biofilms<sup>52,53</sup>. |
| Young's modulus of EPS envelope (Hertz) | $E_{\text{EPS}}$ | [0.001–0.215] | kPa | Varied. |
| Poisson's ratio of EPS envelope (Hertz) | $\nu_{\text{EPS}}$ | 0.45 | – | Within the typical range for biofilms<sup>52,53</sup>. |
| Internal viscosity | $\eta_{\text{int}}$ | 100 | Pa s | Optimized for stability without altering emergent structural response. |
| EPS equilibrium thickness | $L_{\text{EPS}}$ | [300–700] | nm | Varied based on the measured thicknesses (Fig. 1f). |
| Steric repulsive pressure (AdG) | $p_0$ | [0.01–10] | Pa | Varied. |
| *Friction* |  |  |  |  |
| Medium drag coefficient | $\eta^d$ | $7.5 \times 10^{-3}$ | $\text{nNs}\mu\text{m}^{-1}$ | Optimized for stability without altering emergent structural response. |
| Newborn friction coefficient | $\xi_{\text{nb}}$ | 5.0 | $\text{Ns m}^{-1}$ | Optimized for stability allowing relaxation between particles after binary splitting without discrete positional jumps. |
| Characteristic decay time of newborn friction | $\tau_{\text{nb}}$ | 300 | s | Optimized for stability allowing slow relaxation between particles after binary splitting. |
| Wet contact friction coefficient | $\xi_{\text{cc}}$ | [0.1–10] | $\text{kPa s}\mu\text{m}^{-1}$ | Varied. |
| *Numerical parameters* |  |  |  |  |
| Time step | $\Delta t$ | 0.015 | s | Optimized for speed/stability. |
| Conjugate gradient tolerance | $\epsilon_{\text{max}}$ | $10^{-4}$ | – | Chosen to balance speed with accuracy. |
| *Diffusion model* |  |  |  |  |
| Diffusivity | $D$ | 33 | $\mu\text{m}^2\text{s}^{-1}$ | Previously measured for diffusivity of tobramycin in solution<sup>54</sup>. |
| Bacterial sink strength | $a_s$ | [0.01–1.78] | $\mu\text{m}$ | Varied between single-cell scale and domain-size. |
| Grid resolution | $L_{\text{grid}}$ | 0.75 | $\mu\text{m}$ | Approximate size of a bacterium. |
| Domain height | $H_{\text{grid}}$ | $1.5 H_{\text{max}}$ | $\mu\text{m}$ | Set to $1.5\times$ the maximum biofilm height, leaving sufficient space above the biofilm for the concentration profile to relax. |
| Time step | $\Delta t$ | 0.01 | s | Optimized for speed/stability. |

Parameters specifically for the Hertz model or the Alexander-de Gennes (AdG) model are denoted between brackets.

### 17 Reaction-diffusion model

To simulate antimicrobial treatment of biofilm architectures, we simulate a time-dependent reaction-diffusion equation for an antimicrobial compound with concentration *C* in the presence of point sinks representing the consumption of individual bacteria,

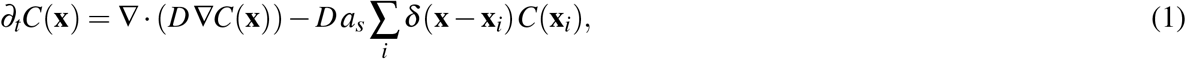

where *D* is the diffusivity of the compound, which we consider homogeneous and isotropic throughout the volume, *a*_*s*_ is the point-sink diffusive size, *δ* is the Dirac delta-function, and **x**_*i*_ are the positions of the sinks placed at the bacterial positions obtained from segmented images. The equation is solved by using a forward time-centered space scheme with a regular rectilinear grid until steady-state is reached over the entire three-dimensional domain. At the top of the domain, a Dirichlet boundary condition is applied with a fixed concentration of *C*_∞_, whereas the other domain interfaces are considered reflective boundaries (∇*C* | _boundary_ = 0). To evaluate the micro-environment of each bacterium, the concentration is then probed at bacterial locations.

## Supplementary Information

### S1 Individual-cell model

Bacteria were modeled as spherocylinders where each spherocylinder *A* = (*i, j*) consists of two nodes with positions **x**_*i*_ and **x** _*j*_ in the centers of both hemispherical ends and a constant radius *R*. The length *L* is determined by the norm of the node positions *L* = ∥**x** _*j*_ − **x**_*i*_∥ and axial direction 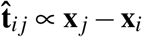. Additionally, the EPS envelope is considered as an additional spherocylinder consisting of the same nodes, thus equal length as *L* but with an increased radius *R* + *L*_EPS_. Nodes belonging to the same spherocylinder have viscoelastic interactions,

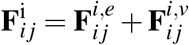

where the elastic force is

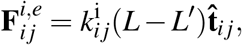

with internal stiffness 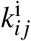, and rest length *L*^′^ and the viscous contribution is given by

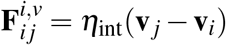

where *η*_int_ is the internal damping coefficient.

#### S1.1 Growth

To model bacterial growth, the rest length *L*^′^ of the elastic interaction between the nodes of the sphero-cylinder is linearly increased every timestep as

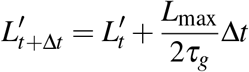

with Δ*t* the time step of the simulation, *L*_max_ the maximum length of a bacteria at which it instigates division, and *τ*_*g*_ the time between subsequent bacterial divisions.

#### S1.2 EPS-dependent repulsion through the Hertz model

We model the interaction between EPS envelopes of contacting spherocylinders through the Hertz model^43^. The steric repulsive force 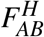 between contacting particles *A* and *B* is

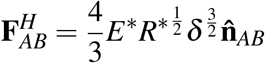

where 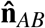 is the normal vector perpendicular to contact plane, *E* is effective combined Young’s modulus of both cells

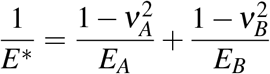

and effective combined radius *R*^*^

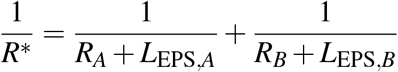

and *δ*_*AB*_ the overlap between contacting particles A,B.

To evaluate the dependency of our results on the exact nature of the formulation of cell-cell interactions, we also modeled repulsive contact through Alexander-de Gennes (AdG) model for contact between polymer brushes^44,45^. The description of this model is provided in SI section S2 and associated results in Supplementary Figure S4.

#### S1.3 Wet contact friction

In addition to steric repulsion, we model viscous interactions between interacting polymer brushes as an area-dependent viscous contact force 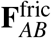 as

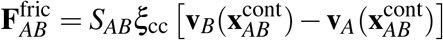

where *S*_*AB*_ is the contact area, *ξ*_cc_ is the contact friction coefficient, and 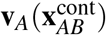 is the velocity of capsule *A* = (*i, j*) at contact point 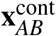 linearly interpolated as

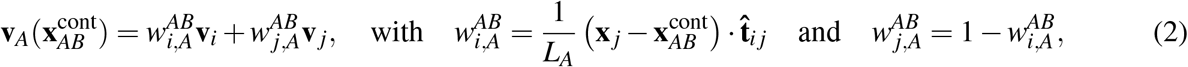

where 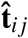 is normal axial vector from *i* to *j* and *L*_*A*_ = ∥**x** _*j*_− **x**_*i*_∥ is the length of the central axis. Velocity at the contact point 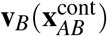 and corresponding weights 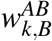 are also analogously defined for capsule *B*.

The resulting drag force is distributed to node *i* of capsule *A* for contact *AB* with capsule *B* as

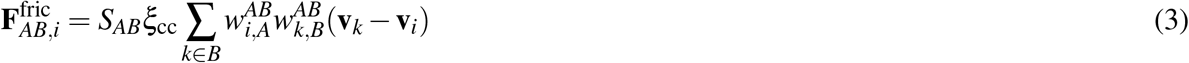

determined by contact area *S*_*AB*_, friction coefficient *ξ*_cc_ and weights *w*^*AB*^ defined in (2). To ensure the stability of the model in case of contacts between one of the spherical caps of the spherocylinder, the weights are capped between 0 and 1.

#### S1.4 Division

When the major axis length of the spherocylinder *L*_*A*_ reaches the maximum length *L*_max_, the spherocylinder *A* = (*i, j*) is split into 2 new separate spherocylinders *B* = (*i, k*) and *C* = (*l, j*) with *i* and *j* the node positions of the original spherocylinder *A* and *k* and *l* at positions

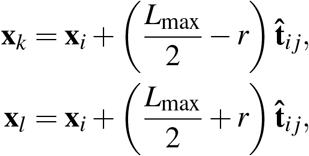

with unit axial vector 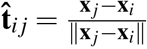.

Since this could lead to potentially large instantaneous overlaps in EPS envelope between newborn spherocylinders *B* and *C*, we add damping that allows these overlaps to gradually relax

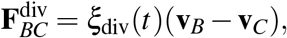

which is resolved to the respective nodes of both newborn capsules *B* and *C* identical to wet contact friction (Eq. 3) and weights (Eq. 2). The friction coefficient *ξ*_div_ decays exponentially over time

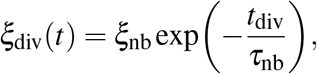

with *t*_div_ the time after division, *ξ*_nb_ the friction coefficient of two newborn spherocylinders, and *τ*_nb_ the characteristic exponential decay time of the friction between newborns. The additional viscous division force **F**^div^ is removed when *ξ*_div_(*t*) < 10^−4^ *ξ*_nb_. This transient viscous force is introduced to relax the initially large overlaps between the EPS envelopes of the two daughter capsules following division. By gradually dissipating these overlaps, the daughter capsules can slowly relax toward their equilibrium positions rather than generating large instantaneous repulsive forces.

#### S1.5 Medium damping

A general isotropic drag force is applied to all nodes,

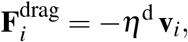

where *η*^d^ is the isotropic drag coefficient. This isotropic drag force does not take into account the specific geometry of the spherocylinders, however, it is relatively small compared to other dissipative forces and it mainly contributes to the numerical stability of the simulations.

#### S1.6 Equation of motion

As bacteria conventionally operate at low Reynolds numbers, we neglect inertial forces. Based on the aforementioned contributions, the force balance for node *i* is

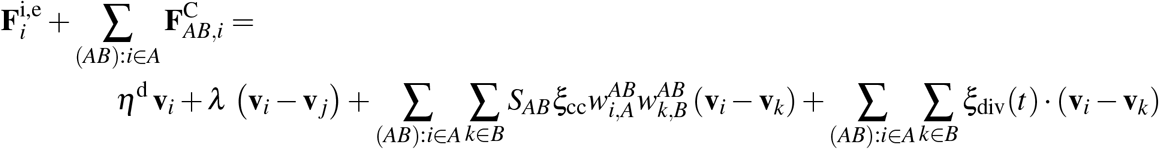

where **F**^C^ constitutes the contact forces (i.e. either Hertz or AdG model). This force balance can be further abbreviated for the full simulation as

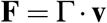

where **F** is the total sum of forces on the nodes and Γ is the combined resistance matrix including off-diagonal contributions for viscous bacteria-bacteria and bacteria-medium interactions. The equation of motion is solved for the node velocities **v** using the conjugate gradient method each iteration. Subsequently, positions of the particles are updated using a semi-implicit Euler integration scheme

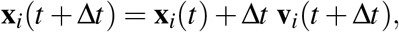

with **v**_*i*_(*t* + Δ*t*) are the projected new velocities obtained at time *t*.

### S2 EPS-dependent repulsion through the Alexander-de Gennes model

For the Alexander-de Gennes model, the EPS envelope is modeled as a brush layer of polymers on the surface of the bacteria. The steric repulsive pressure 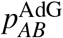 is modeled as two non-interacting polymer brushes surrounding particles *A* and *B* as

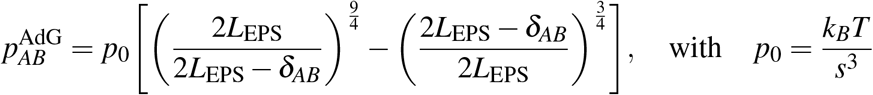

where *k*_*B*_ is Boltzmann’s constant, *T* is the absolute temperature, *L*_EPS_ the equilibrium thickness of polymer brush, *s* the grafting distance between polymers on the bacterial surface, and *δ*_*AB*_ the overlap between the polymer brushes of spherocylinder *A* and *B*, which is obtained by the shortest segment **n**_*AB*_ connecting the segments defined by the nodes of both respective capsules. The first term signifies the increased osmotic pressure due to compression of both polymer brushes, whereas the second term accounts for the decrease in elastic energy. The effective steric interaction force 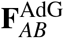 is given by

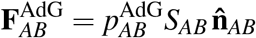

where the contact area is given by 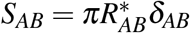 with effective particle radius R* similarly defined as in the Hertz model and 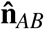 the normal vector perpendicular to the contact plane. For computational stability, 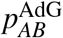 was limited to 100 *p*_0_ in the case of very high overlaps.

### S3 Spatial killing heterogeneity derivation

We want to characterize the influence of the micro-architecture of the resulting colony on the killing probability

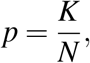

where *K* is the number of dead cells and *N* is the total number of cells. In particular, we are interested in the difference between the periphery of the colony, characterized by low density, and the core of the colony, characterized by high density.

We can capture the heterogeneity of the killing probability by the variance of the mean killing probability *p*(*ρ*) at a given density *ρ*,

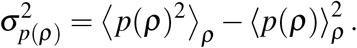

If the killing probability is density independent, this variance vanishes. In the case where all cells in the periphery are killed, while all cells occupying the core are preserved, the variance reduces to

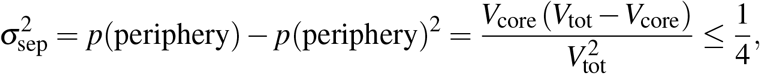

where *V*_core_ is the volume occupied by the colony core and *V*_tot_ is the total volume of the colony, and the upper bound is reached when the core’s and periphery’s volumes are equal. Note that this inequality holds even in the general case, so the variance of the mean killing rate is bounded by

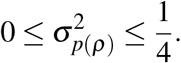

In practical settings, due to the discrete nature of the bacterial colony, there is a dependency between the density profile and the spatial resolution. To investigate the influence of scale, we extend the variance of the mean killing rate to explicitly contain the scale *l* as

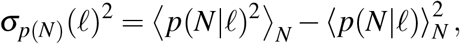

where *N* is the number of cells inside a box of size *l*, and the mean killing probability *p*(*N*|*l*) is defined as

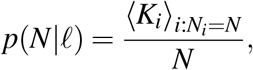

where the mean value is taken over all boxes containing exactly *N* bacteria. Note that the density is linearly proportional to the total number of cells in a given box, *ρ* = *N/l*^3^. The boxes are uniformly sampled over the whole domain.

For scales much smaller than the typical spacing between bacteria, a single box contains at most one cell, making the whole population effectively homogeneous. Similarly, for a box spanning the entire domain, we again obtain a homogeneous population, and the variance should again vanish.

### S4 Bulk slab approximation for reaction-diffusion

In the limit of homogeneously distributed sinks over a domain with volume *V*, equation (1) simplifies to

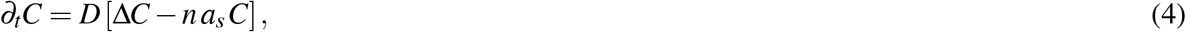

where *n* = *N/V* is a volumetric sink concentration in the domain. Furthermore, if the biofilm can be considered as a homogeneous slab with height *H*, we can find explicitly the steady-state solution of (4) as

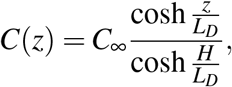

where the diffusive length-scale *L*_*D*_ emerges as

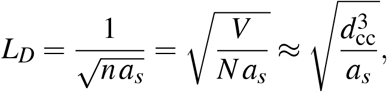

with typical inter-cell spacing *d*_cc_. The expected mean value over the slab is then

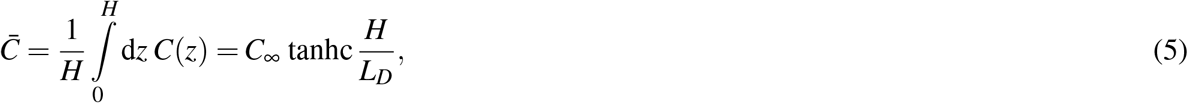

where tanhc *x* = tanh *x/x*. And the variance over the profile is then given by

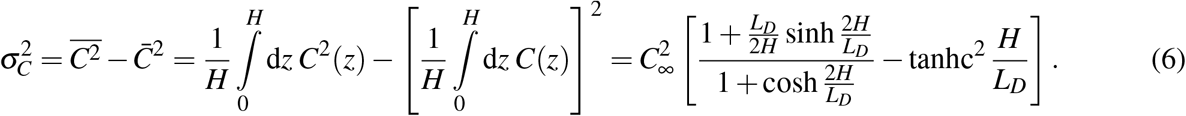

Note that when the diffusive length-scale is much larger than the slab height (*L*_*D*_ ≫ *H*), the average concentration is close to the boundary condition 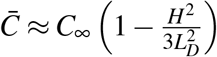 while the variance scales the relative diffusive length scale as

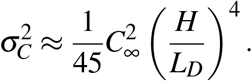

Conversely, when the sinks dominate (*L*_*D*_ ≪ *H*), the concentration decreases as

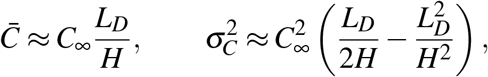

while variance has a clear maximum. Eliminating *L*_*D*_*/H* using 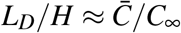 gives the second-order relation

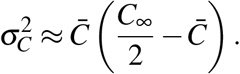

Introducing the normalized mean concentration 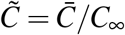 therefore yields

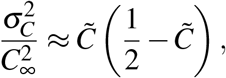

which is maximal for

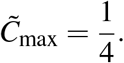

We also found that the exact numerical solution of (6) with respect to (5) is 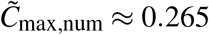 which is quite close to this value. Thus, the occurrence of maximal concentration heterogeneity near 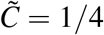 follows directly from the sink-dominated slab solution.

### S5 Isolated sink regime for reaction-diffusion

For a set of isolated sinks, the steady-state solution of (1) can be approximated by the set of concentration profiles *C*_*i*_ obtained as the solution for a single sink

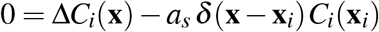

with boundary condition *C*_*i*_(**x**) → *C*_∞_ for ∥**x** − **x**_*i*_∥ → ∞. In 3D, the steady state solution of a point sink is given by

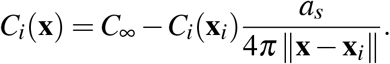

The concentration profile diverges at the sink position, where we evaluate the antimicrobial concentration experienced by the bacteria. This indicates that the point-sink approximation is no longer suitable. To rectify this, we assume that consumption is uniform over the cell volume (*V*_*i*_), allowing us to determine the average concentration 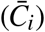 experienced by the bacterium *i* at its location (**x**_*i*_).

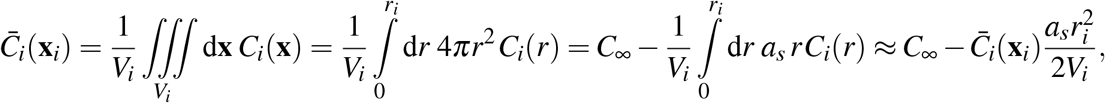

where we, for simplicity, assume the cell is spherical, and also in the last step, we approximated the concentration profile with the value in the middle. From there, the mean concentration inside the cell can be obtained as

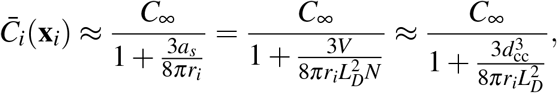

which, in the limit of large separation (*d*_cc_ ≫ *L*_*D*_ and *L*_cc_ ≫ *r*_*i*_), scales as

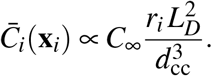

## 2 Supplemental figures

**Supplementary Fig. 1:**
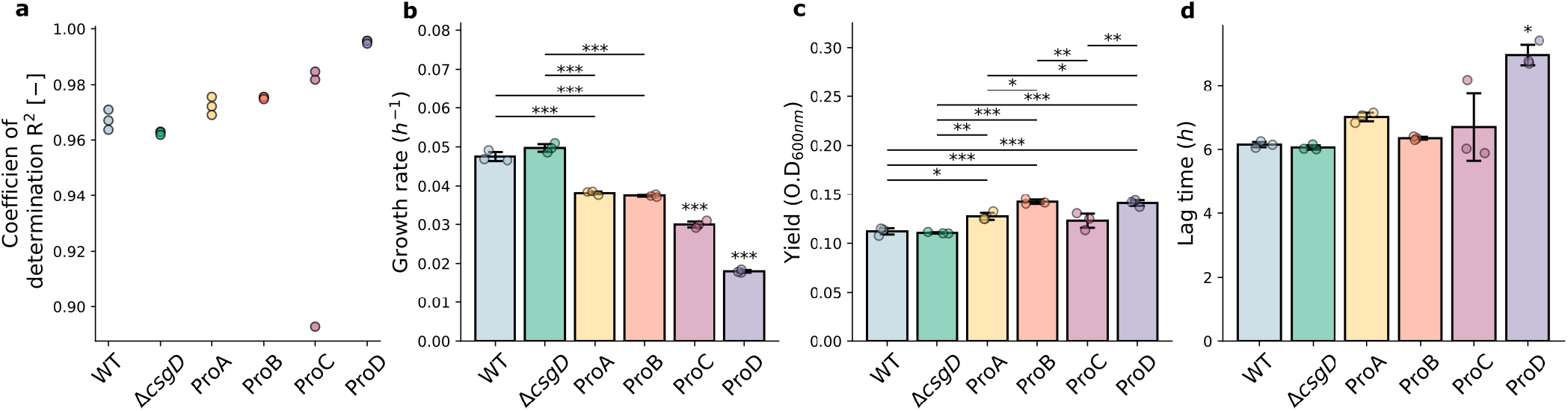
Parameter extraction from Gompertz fits to planktonic growth curves. **a**, Coefficient of determination (*R*^2^) as a measure of goodness of fit. **b**, Extracted instantaneous growth rate *µ*. **c**, Extracted yield *A* after 24 h of planktonic growth. **d**, Lag time *λ* of the different strains. Bars indicate means ± standard deviation of *n* = 3 independent replicates.

**Supplementary Fig. 2:**
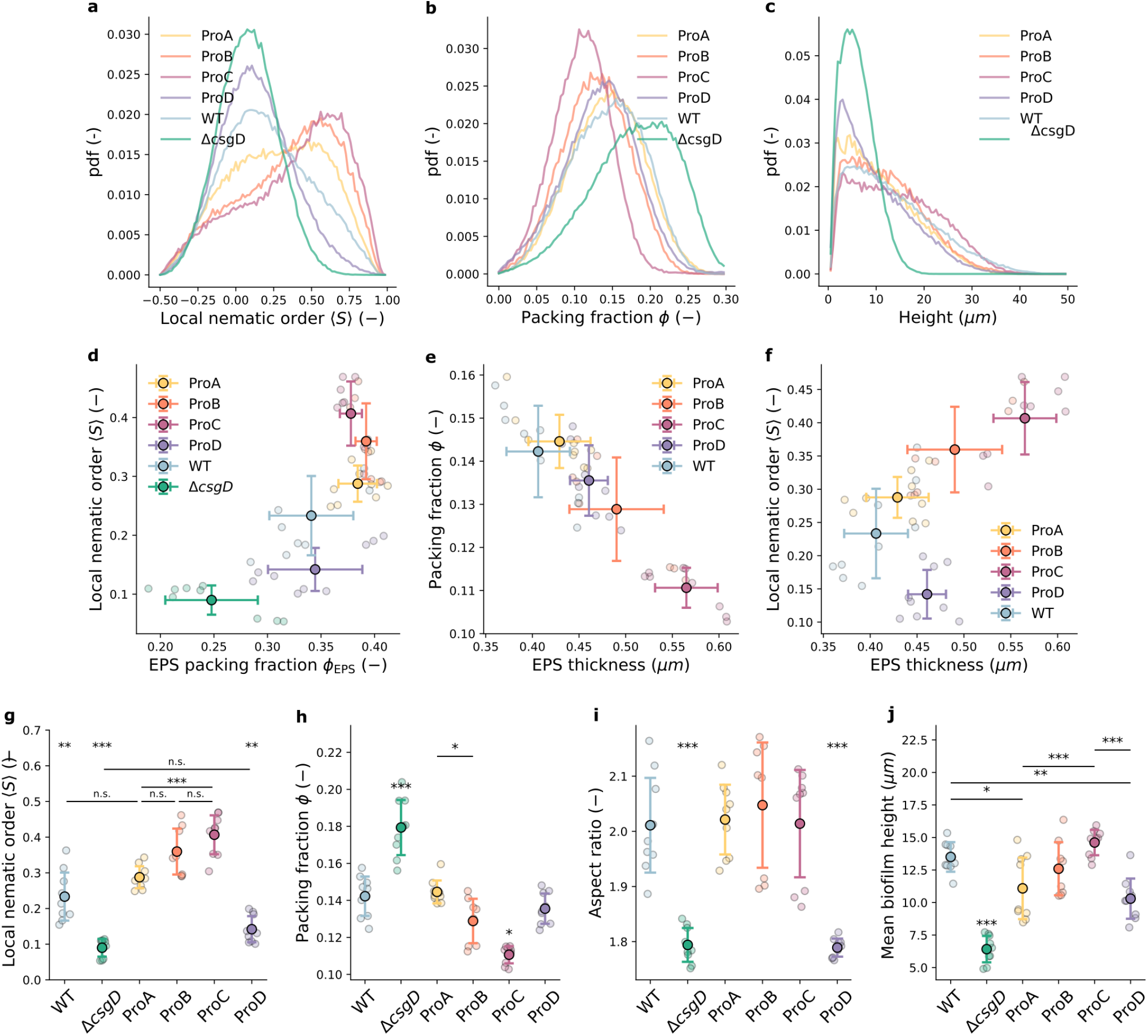
Architectural characterization of biofilms with increasing EPS production. **a–c**, Distributions of cell-level local nematic order (**a**), packing fraction (**b**), and vertical position (**c**) across independent biofilm replicates. **d–f**, Architectural scaling of local nematic order as a function of EPS packing fraction (**d**), packing fraction as a function of EPS thickness (**e**), and local nematic order as a function of EPS thickness (**f**). **g–j**, Comparison of biofilm architectural parameters across strains, with points and error bars representing the mean and standard deviation across independent biofilm replicates. Means and standard deviations of *n* = 9 independent biofilms are shown. Statistically significant pairwise comparisons were determined using one-way analysis of variance (ANOVA) followed by Tukey’s honestly significant difference (HSD) test. ^***^*P* < 0.001; ^**^*P* < 0.01; ^*^*P* < 0.05; n.s., not significant.

**Supplementary Fig. 3:**
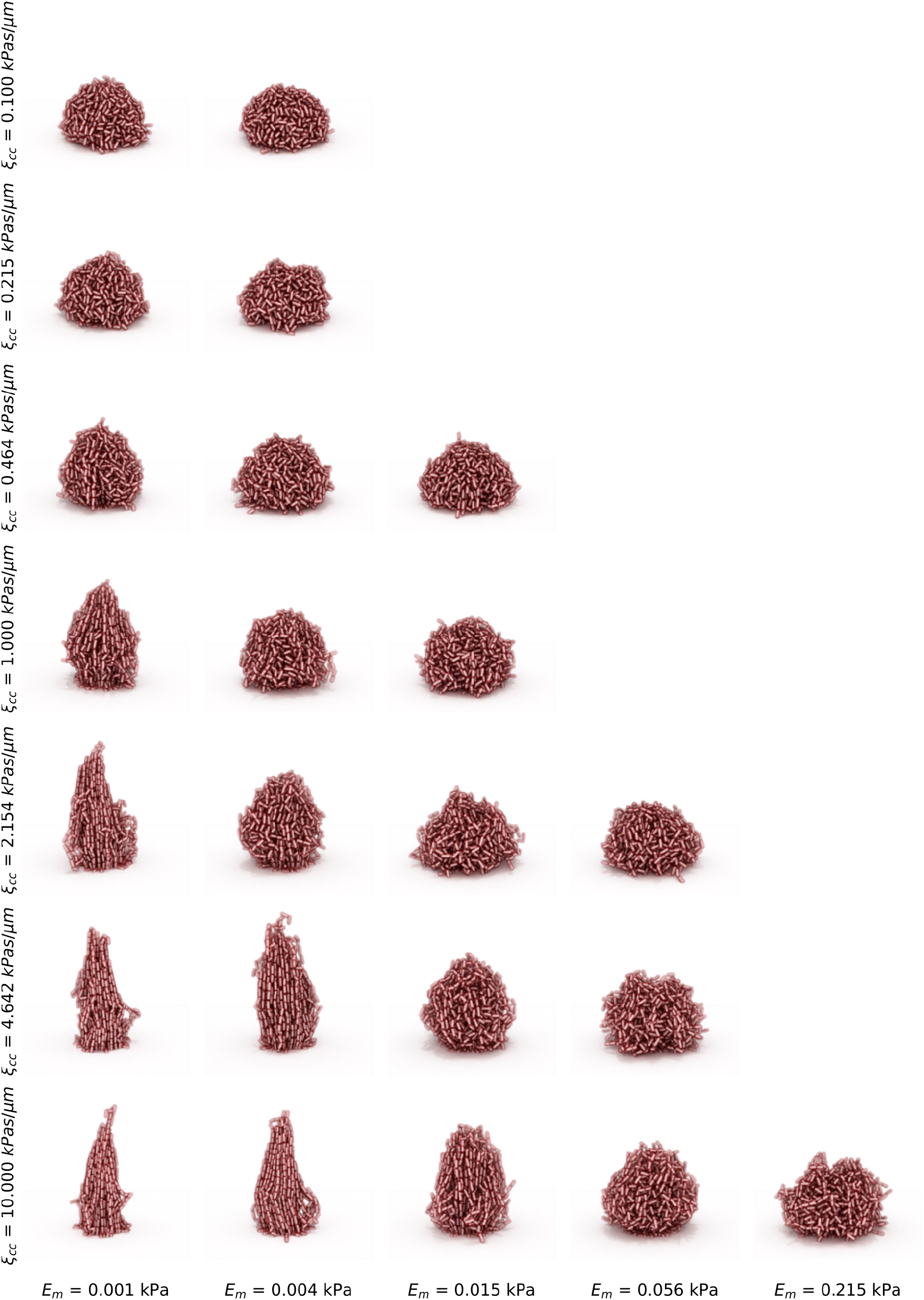
Model visualization with fixed EPS thickness *L*_EPS_. Visualization of the simulation endpoint as a function of viscous inter-envelope friction *ξ*_cc_ (vertical axis) and envelope stiffness *E*_EPS_, with a fixed EPS thickness of *L*_EPS_ = 500 nm and generation time *τ*_*g*_ = 20 min.

**Supplementary Fig. 4:**
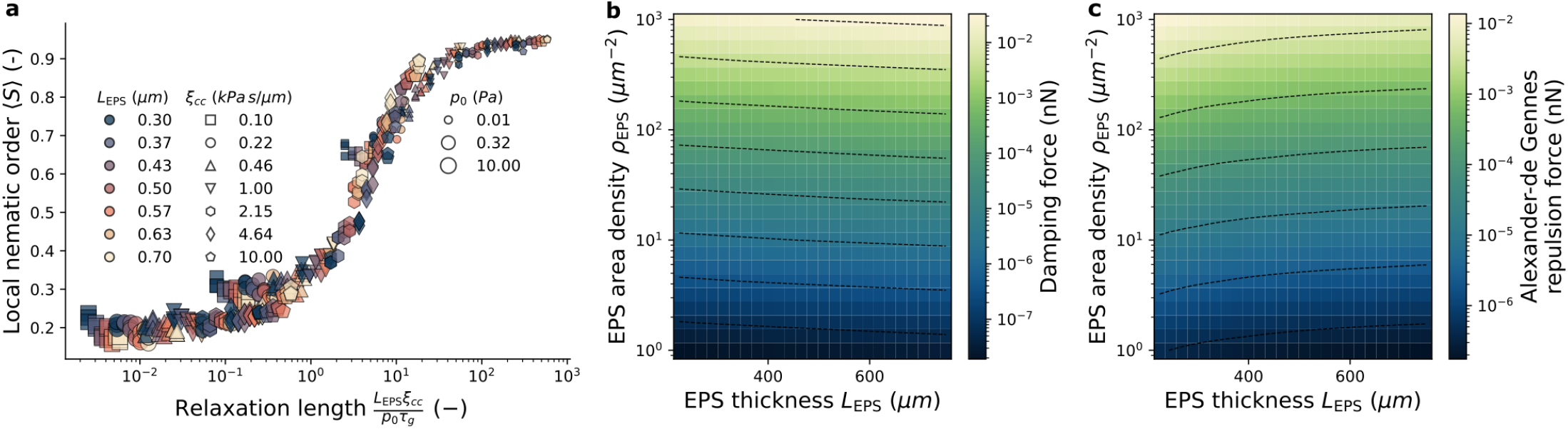
Architectural transitions are largely invariant to the specific mechanical signature of the local cell-cell interactions. **a**, Simulating biofilm growth using the Alexander-de Gennes model rather than a simple Hertzian repulsive force produces qualitatively similar behavior as a function of the relaxation length. This length scale balances the EPS thickness with a hydrodynamic length scale, *ξ*_cc_*/*(*p*_0_*τ*_*g*_), where *ξ*_cc_ is the viscous inter-envelope friction, *p*_0_ is the reference pressure at the cell surface, and *τ*_*g*_ is the generation time. **b**,**c**, Scaling of the viscous damping (**b**) and repulsive Alexander-de Gennes forces (**c**) as a function of microscopic EPS parameters, including the EPS surface density *ρ*_EPS_ and thickness *L*_EPS_.

**Supplementary Fig. 5:**
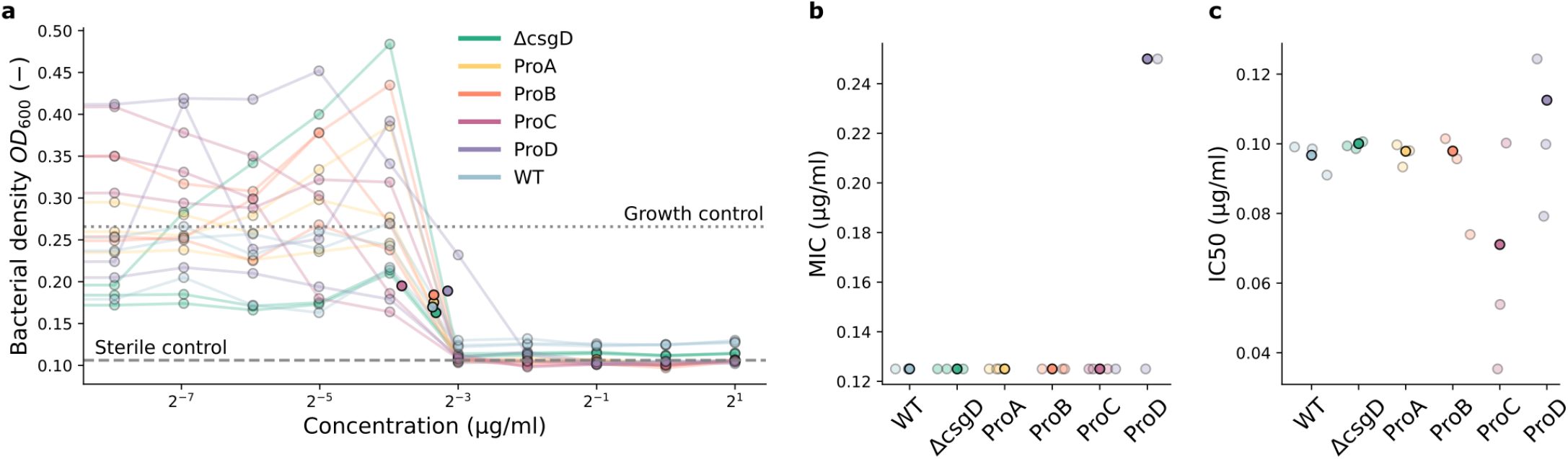
Cefotaxime MIC assay. **a**, Dose-response curve obtained by endpoint measurement of bacterial density as a function of cefotaxime concentration. Dashed and dotted lines indicate the average bacterial density of the sterile control and growth control, respectively, with the growth control corresponding to WT in the absence of CTX. **b**, MIC extracted as the lowest concentration at which measurable bacterial growth was detected, defined as a bacterial density greater than the average of the sterile control plus five standard deviations. **c**, IC50 concentration for all strains obtained from a Hill fit to independent dose-response curves. Solid points indicate the mean of *n* = 3 independent repeats.

**Supplementary Fig. 6:**
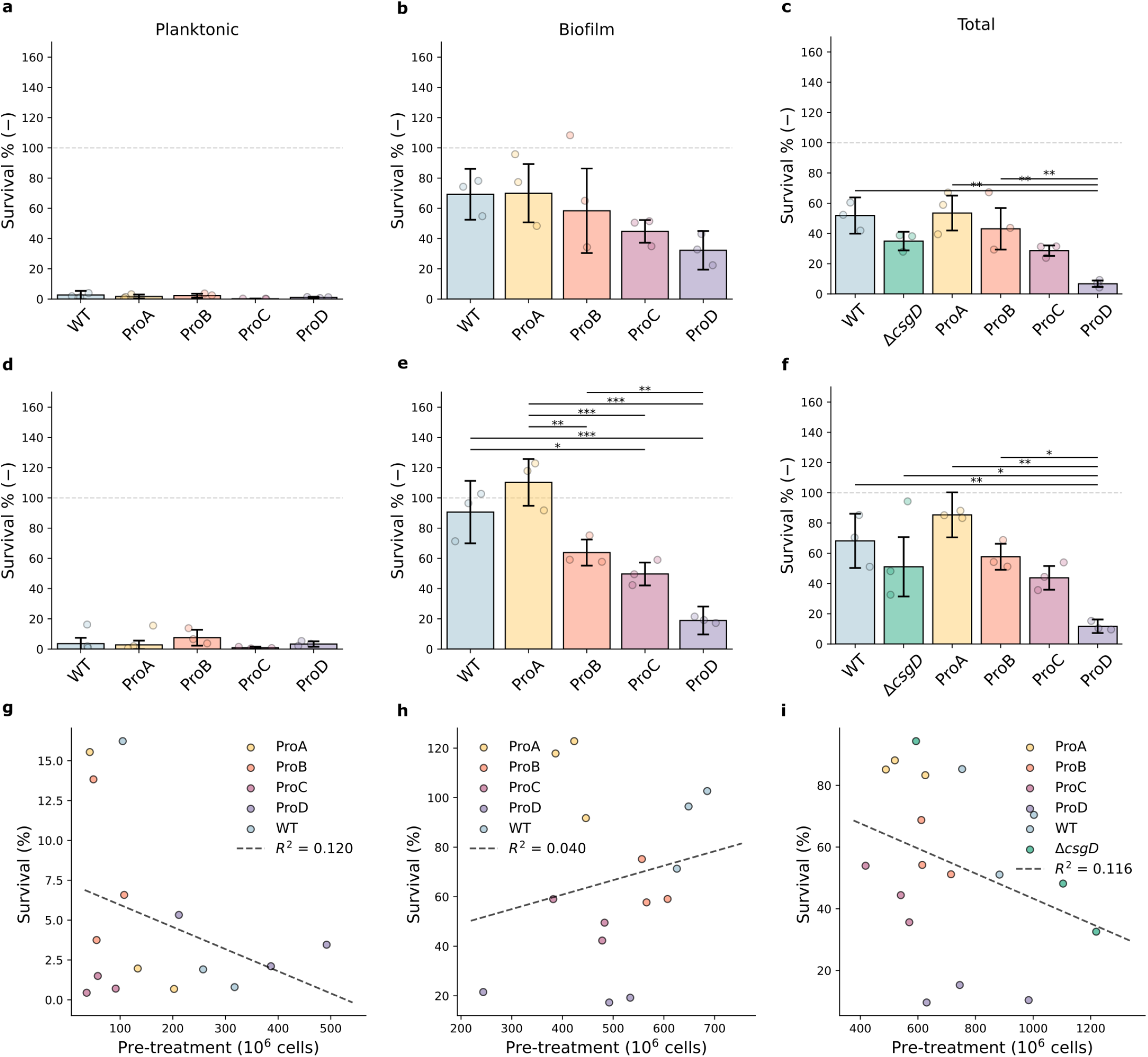
Bulk antimicrobial survival following cefotaxime treatment. **a–c**, Population survival after treatment with 500 *µ*gmL^−1^ cefotaxime for 4 h, compared with an untreated control measured after 24 h + 4 h, for the planktonic (**a**), biofilm (**b**), and total (**c**) populations. **d–f**, Population survival after treatment with 500 *µ*gmL^−1^ cefotaxime for 4 h, compared with the initial biofilm architecture at 24 h, for the planktonic (**d**), biofilm (**e**), and total (**f**) populations. **g–i**, No strong correlation was observed between cell number before treatment and eventual survival for the planktonic (**g**), biofilm (**h**), and total (**i**) populations. Points indicate the average of three technical repeats for each of three independent biological repeats, whereas bars and error bars indicate the mean and standard deviation across the *n* = 3 biological repeats. Statistically significant pairwise comparisons were determined using one-way analysis of variance (ANOVA) followed by Tukey’s honestly significant difference (HSD) test. ^***^*P* < 0.001; ^**^*P* < 0.01; ^*^*P* < 0.05; n.s., not significant.

**Supplementary Fig. 7:**
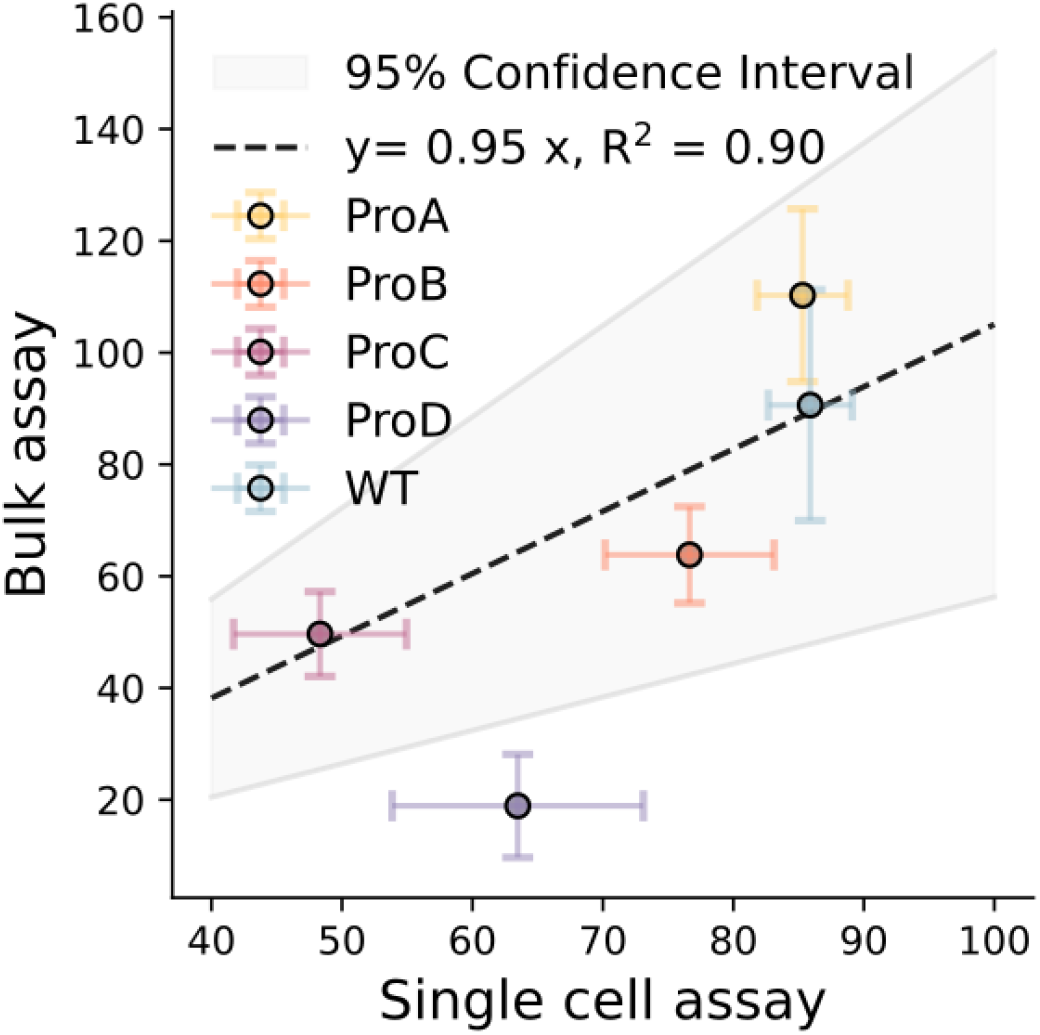
Comparison between population-level and microscopy-based single-cell survival. The bulk assay shows a near one-to-one correlation (slope = 0.95, *R*^2^ = 0.90) with the microscopy-based single-cell assay. Points represent the average values of three independent repeats (*n* = 3), with error bars indicating the standard deviation. The fit was performed using orthogonal distance regression.

**Supplementary Fig. 8:**
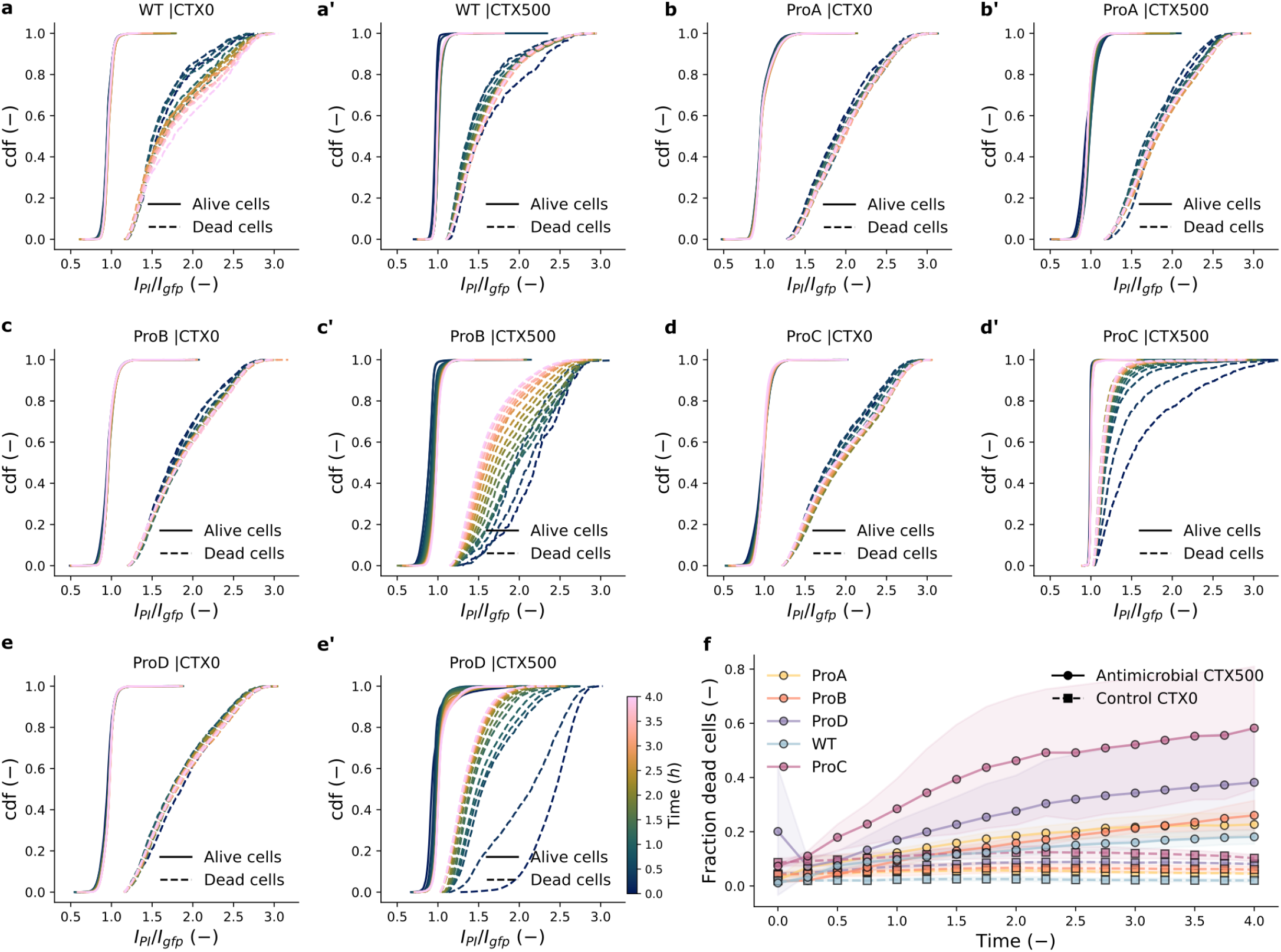
Single-cell classification of cell viability and killing dynamics. **a–e’**, Gating strategy and intensity-ratio distributions (*I*_PI_*/I*_GFP_) of the propidium iodide stain (*I*_PI_) and constitutive GFP signal (*I*_GFP_) used to classify membrane-compromised (dead) and membrane-uncompromised (viable) cells. Profiles are shown for untreated control (CTX0) and antimicrobial-treated (CTX500, 500 *µ*gmL^−1^ cefotaxime, 4 h treatment) biofilms across all strains. **f**, Temporal fraction of membrane-compromised cells over 4 h of treatment (CTX500, solid lines with circles) or control incubation (CTX0, dashed lines with squares) for all five strains.

**Supplementary Fig. 9:**
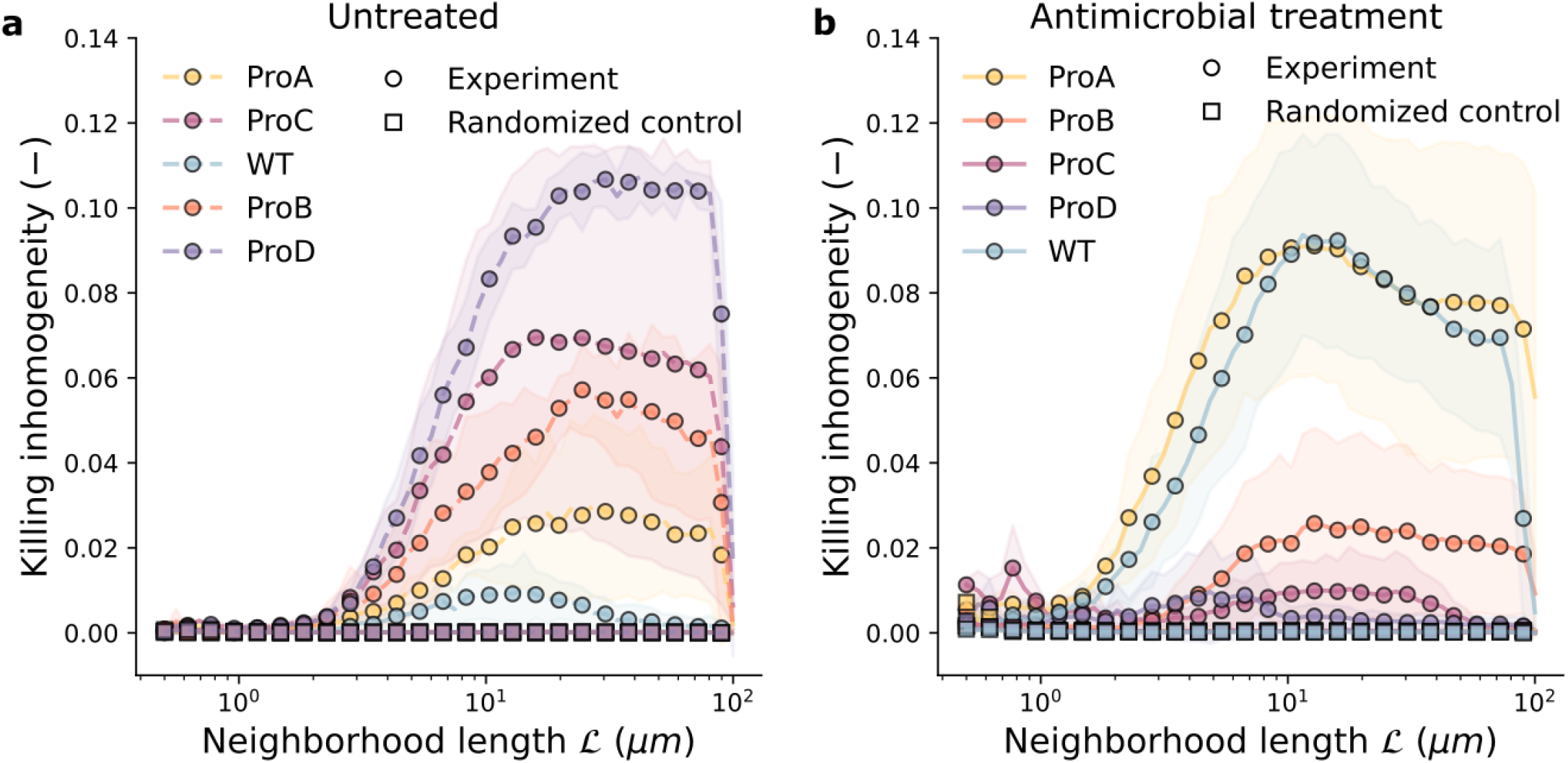
Killing inhomogeneity across neighborhood length scales. **a**, In untreated biofilms, spontaneous cell death is non-randomly distributed, exhibiting a distinct peak in killing inhomogeneity around neighborhood length scales of *l* ≈ 5–10, *µ*m across all strains. **b**, Following antimicrobial treatment, WT and ProA maintain high killing inhomogeneity extending to larger spatial scales. In contrast, high-matrix-producing strains (ProB, ProC, and ProD) display progressively weaker spatial heterogeneity, indicating a more homogeneous spatial distribution of cell death. Experimental curves (solid lines) are plotted alongside randomized controls.

**Supplementary Fig. 10:**
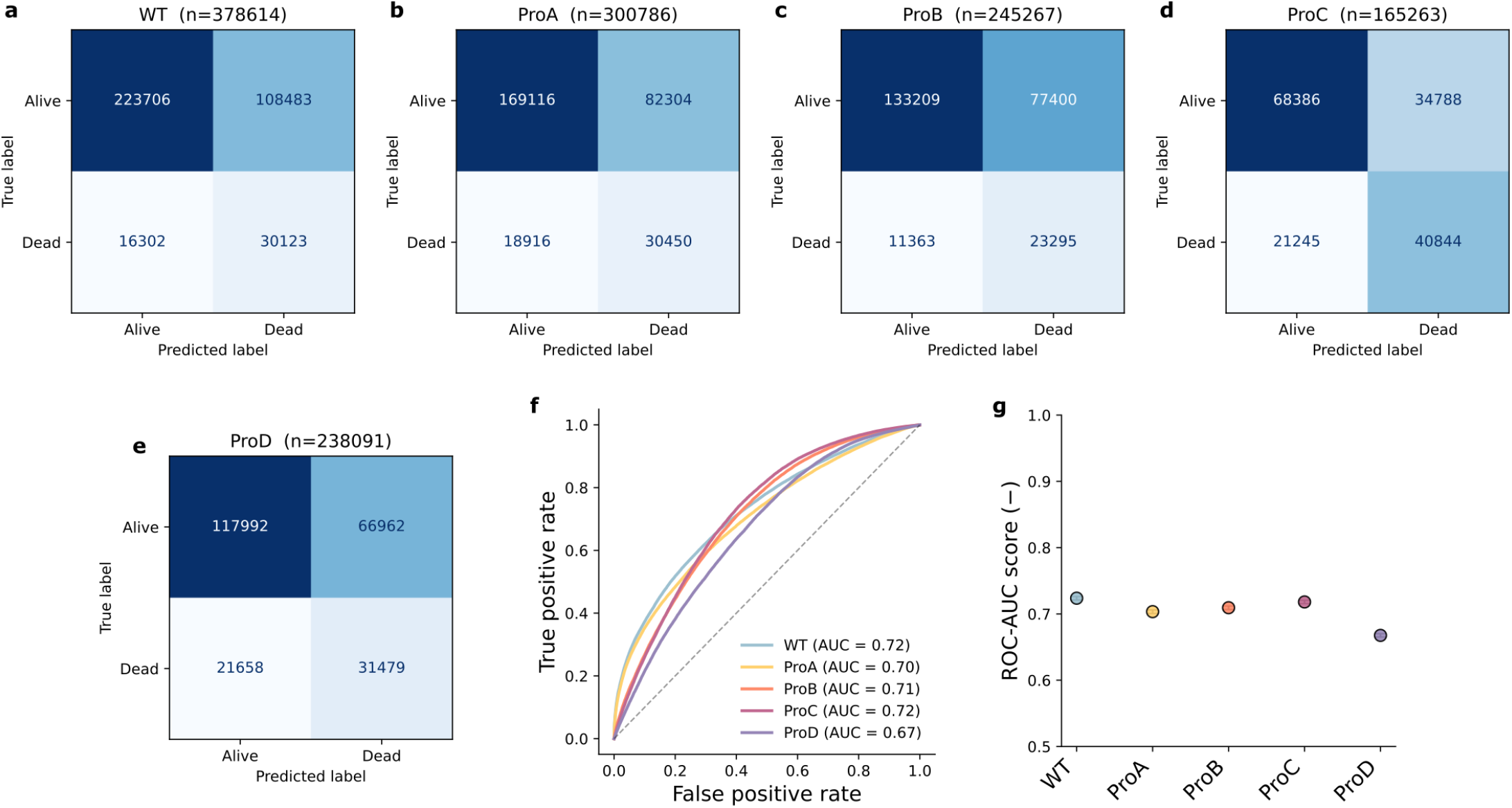
Performance and validation of logistic regression models predicting single-cell death based on biofilm architectures. **a–e**, Confusion matrices showing true and predicted cell states (viable/dead) for all strains, with the total number of evaluated cells (*n*) indicated. **f**, Receiver operating characteristic (ROC) curves showing model predictive accuracy across strains, with the area under the curve (AUC-ROC) ranging from 0.63 to 0.72. **g**, Model classification performance summarized by the mean ROC-AUC score for each strain across cross-validation splits.

**Supplementary Fig. 11:**
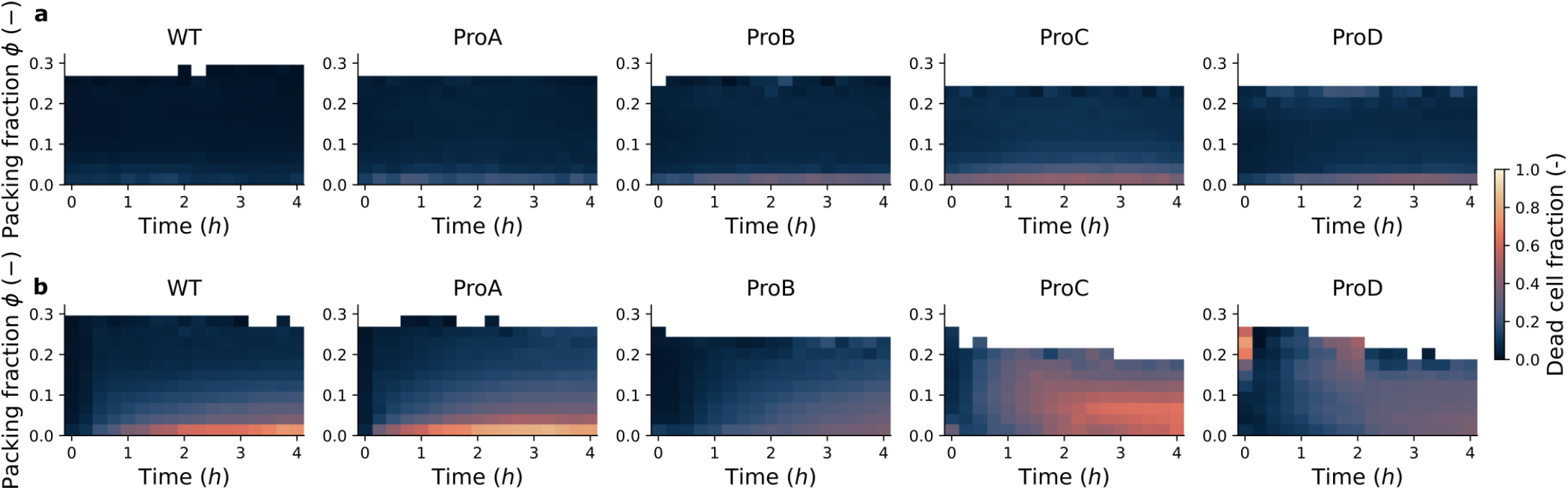
Dead-cell probability as a function of the temporal evolution of local packing fraction. **a**,**b**, Kymographs showing local packing fraction over time for each strain, with color indicating the fraction of dead cells under treated (**a**; 500 *µ*gmL^−1^ cefotaxime, 4 h treatment) and untreated (**b**) conditions. Data represent the average across three independent biofilms (*n* = 3).

**Supplementary Fig. 12:**
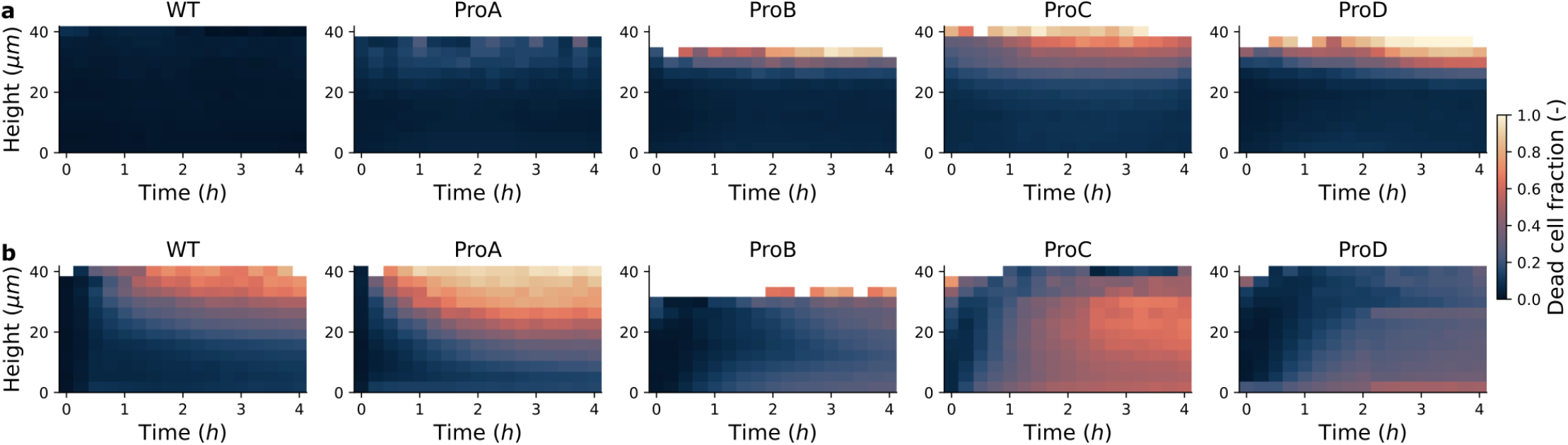
Dead-cell probability as a function of the temporal evolution of vertical cell position. **a**,**b**, Kymographs showing normalized cell height over time for each strain, with color indicating the fraction of dead cells under treated (**a**;500 *µ*gmL^−1^ cefotaxime, 4 h treatment) and untreated (**b**) conditions. Data represent the average across three independent biofilms (*n* = 3).

**Supplementary Fig. 13:**
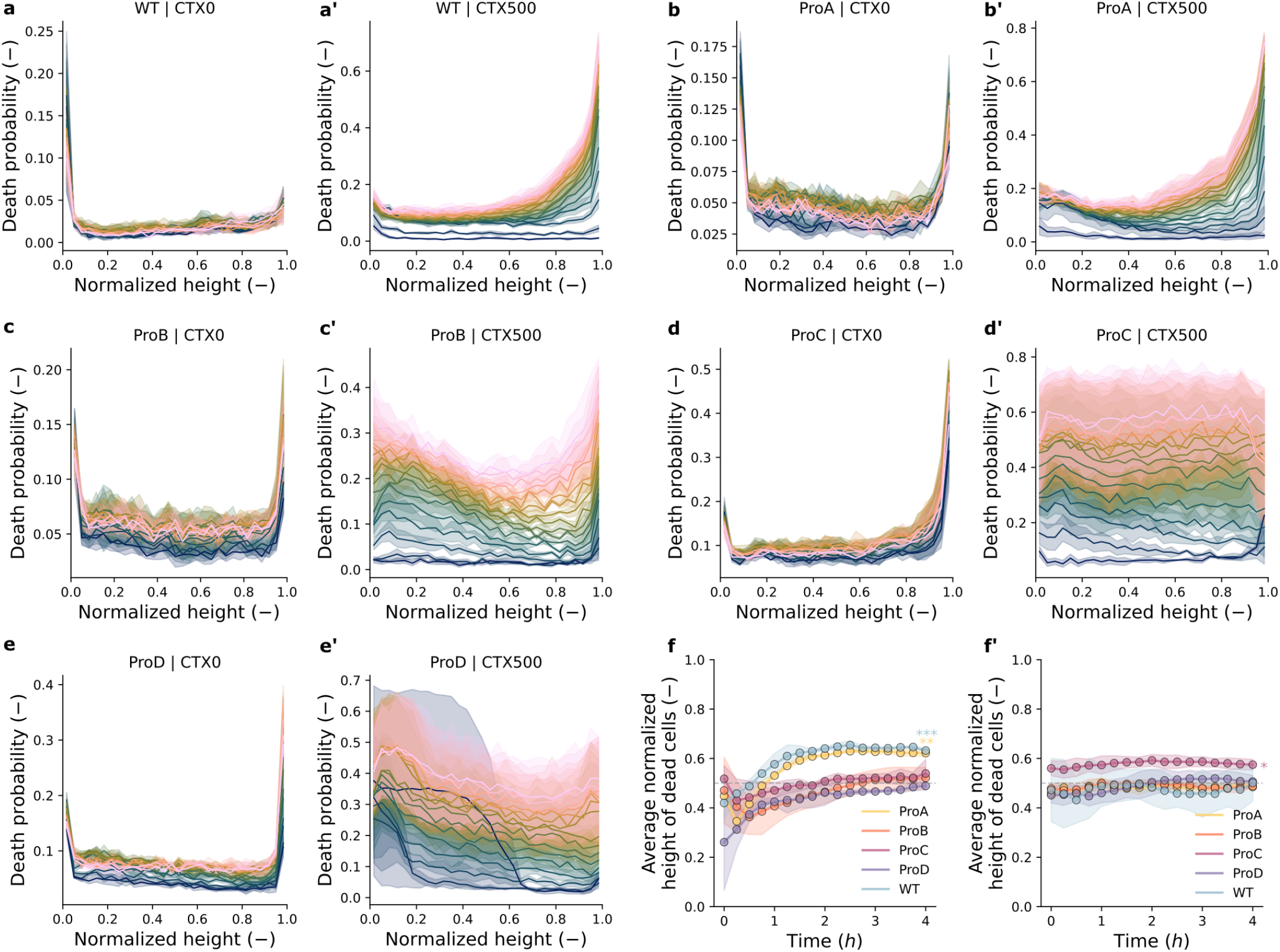
Depth-dependent killing probability profiles across different biofilm strains. **a–e’**, Death probability as a function of normalized biofilm height for untreated control (CTX0) and treated (CTX500) conditions across all strains, illustrating spatial differences between vertically uniform killing (ProB, ProC, and ProD) and depth-dependent killing (ProA and WT). **f**,**f’**, Average normalized height of dead cells plotted over treatment time in the presence (**f**) and absence (**f’**) of antimicrobials, demonstrating that killing remains targeted to the upper layers for WT and ProA, while remaining vertically uniform for ProB, ProC, and ProD. Statistical significance was assessed using a one-sample *t*-test against 0.5, the expected mean height for a vertically uniform distribution. ^***^*P* < 0.001; ^**^*P* < 0.01; ^*^*P* < 0.05.

**Supplementary Fig. 14:**
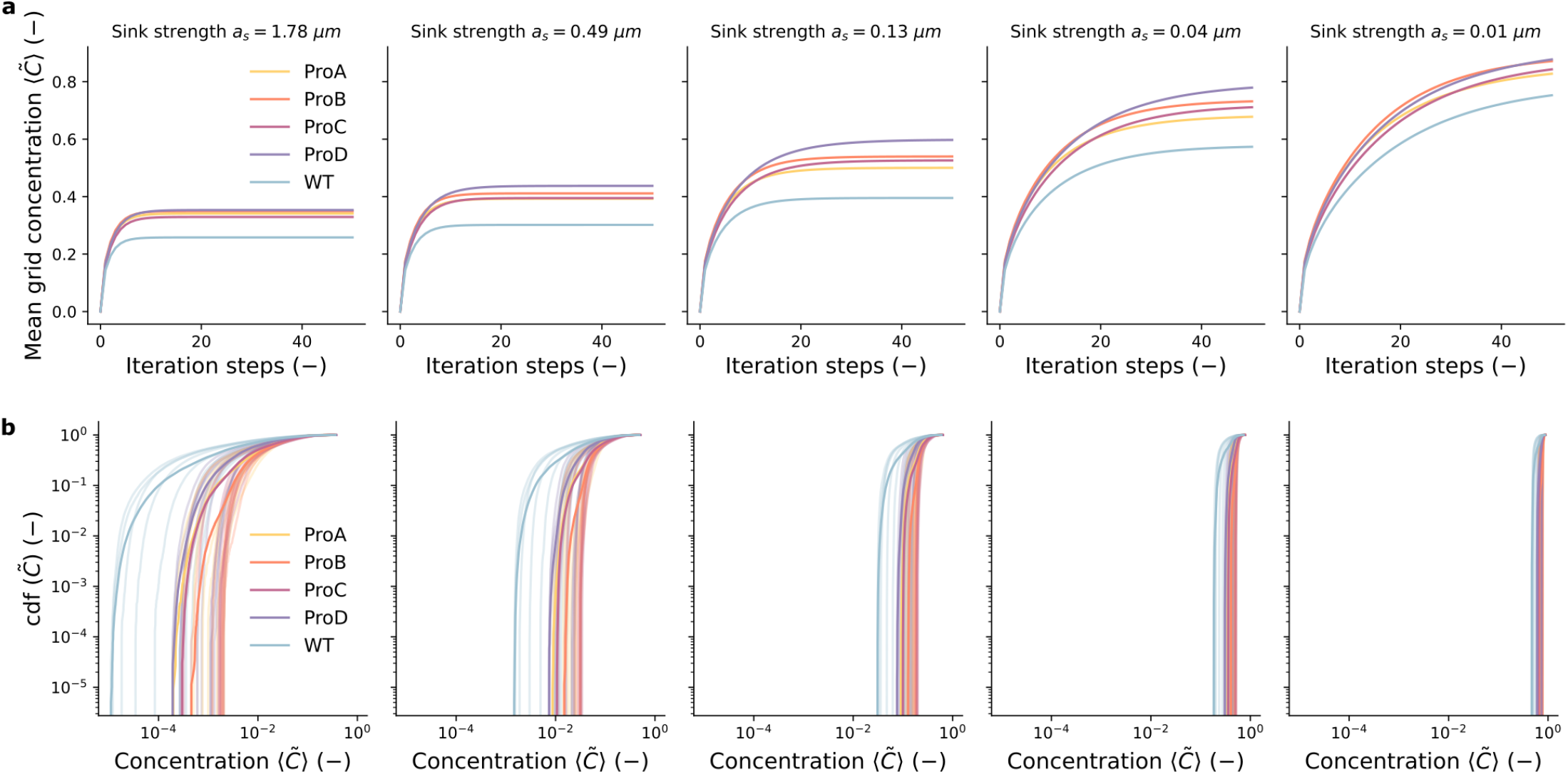
Convergence and distribution of simulated local antimicrobial concentration fields. **a**, Mean grid concentration as a function of simulation iteration steps for increasing point-sink strengths (*a*_*s*_ = *k/D*), demonstrating numerical convergence to steady state across all strain architectures. **b**, Cumulative distribution functions (CDFs) of the normalized local concentration experienced by cells across all strain architectures under different simulated sink strengths (*a*_*s*_). Solid profiles are obtained from simulations on nine independent biofilm architectures (*n* = 9).

**Supplementary Fig. 15:**
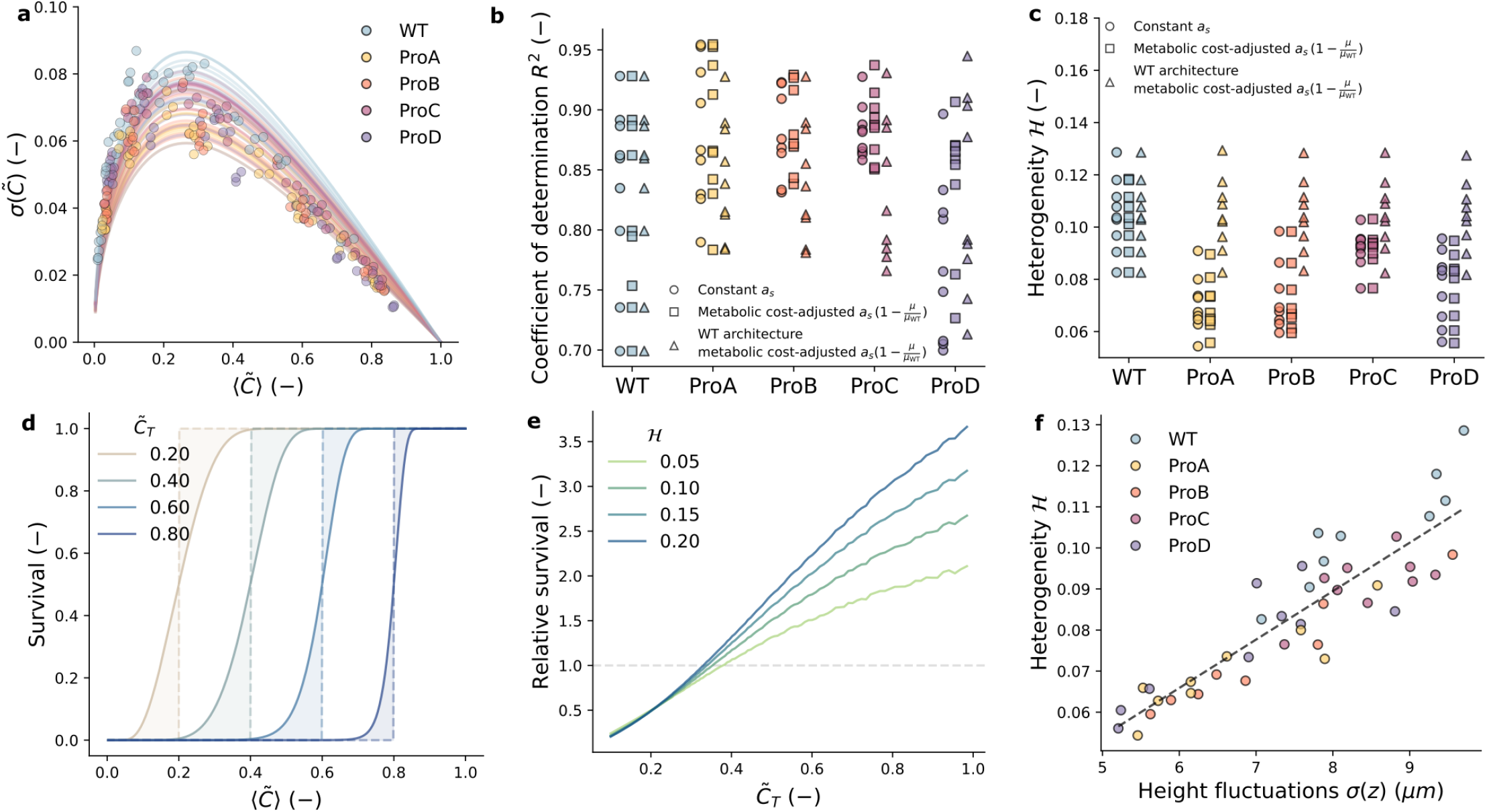
Characterization of concentration-field heterogeneity and its role in survival predictions. **a**, Standard deviation of local concentration *σ* (*C*) plotted against the mean concentration ⟨*C*⟩ for the metabolic cost-adjusted sink strength *a*_*s*_, fitted using a bulk approximation to extract structural heterogeneity *ℋ*. **b**,**c**, Coefficient of determination (*R*^2^) (**b**) and inferred architectural heterogeneity *ℋ* (**c**) for different transport models: constant *a*_*s*_, metabolic cost-adjusted *a*_*s*_(1 −*µ/µ*_WT_), and metabolic cost-adjusted *a*_*s*_(1 − *µ/µ*_WT_) combined with WT architecture. **d**, Theoretical survival fraction as a function of mean concentration for heterogeneous structures (*ℋ* = 0.10, solid lines) and perfectly uniform architectures (*ℋ* = 0.0, dashed lines), demonstrating how structural heterogeneity provides a survival advantage at higher killing thresholds (bulk regimes). **e**, Relative survival plotted against the normalized killing threshold 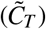 relative to perfectly uniform structures. **f**, Architectural heterogeneity *H* scales linearly (slope = 0.012, *R*^2^ = 0.79) with vertical biofilm height fluctuations *σ* (*z*). Data are obtained from nine independent biofilm architectures (*n* = 9).

**Supplementary Fig. 16:**
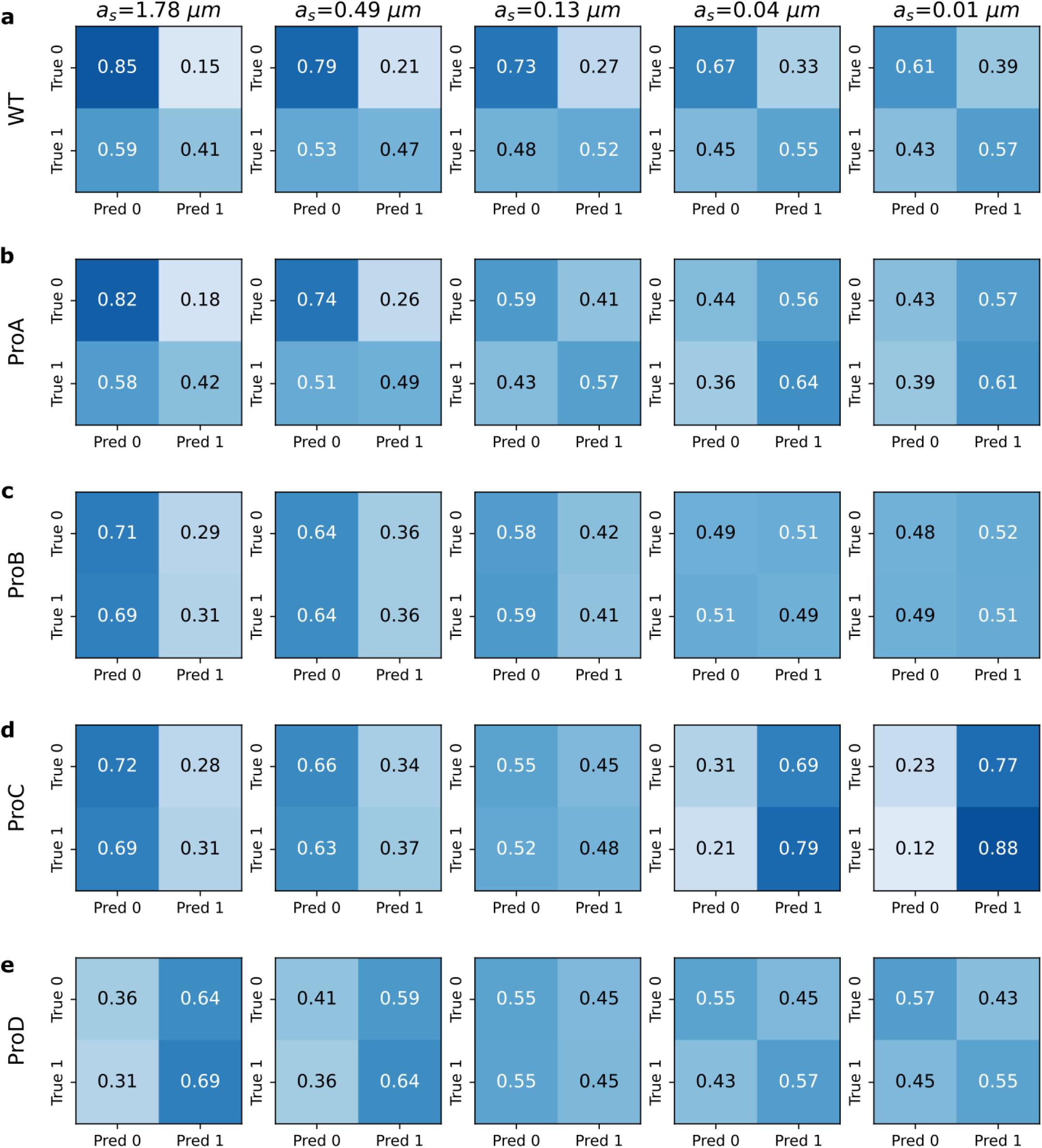
Confusion matrices for single-cell survival predictions based on simulated concentration fields. Confusion matrices showing true and predicted cell states (alive/dead) for all strains evaluated across five simulated point-sink strengths. Data are obtained from three independent biofilm replicates per strain (*n* = 3).

**Supplementary Fig. 17:**
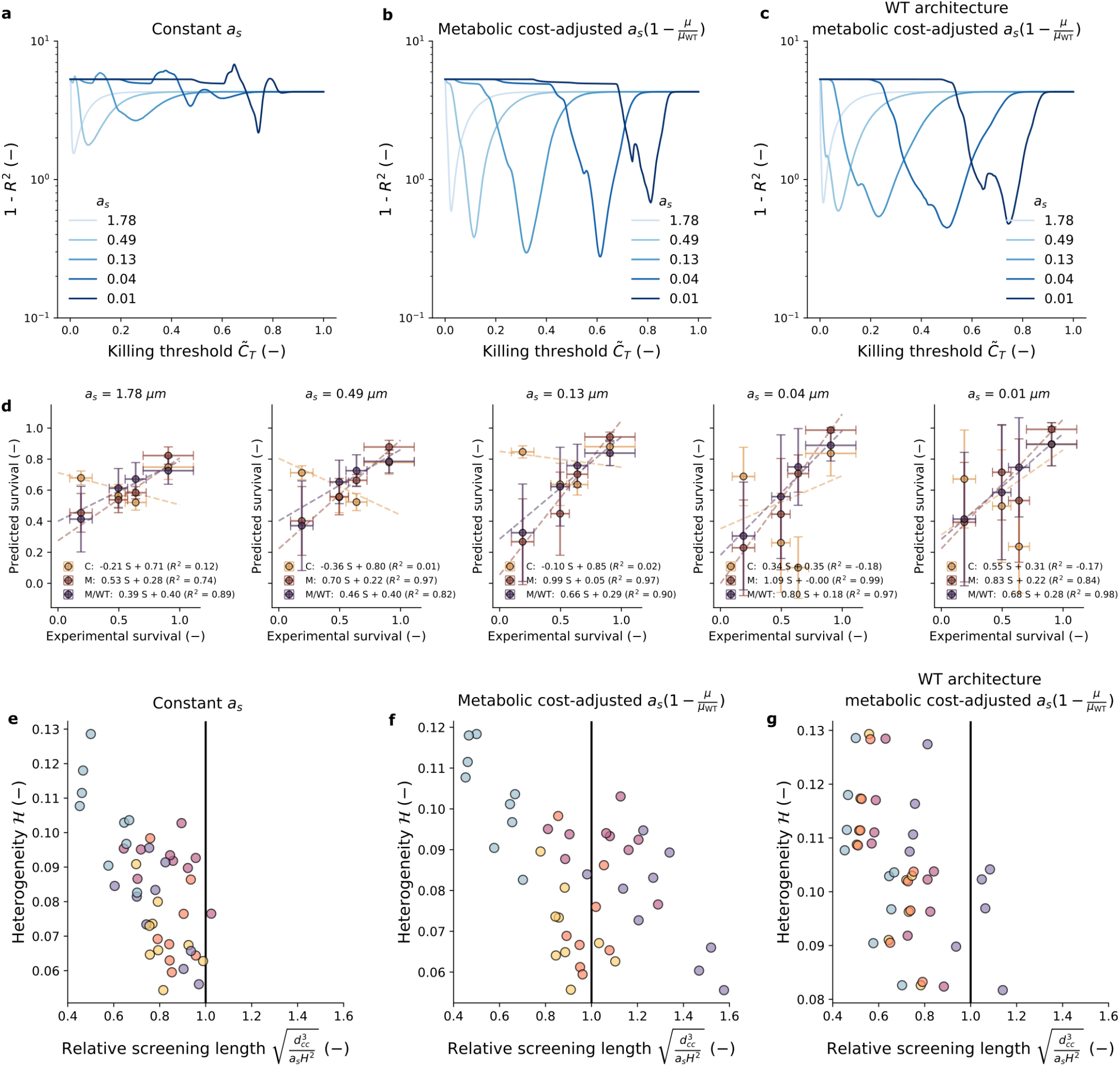
Sensitivity analysis and bulk survival predictions under different transport assumptions. Goodness of fit (1 − *R*^2^) plotted against the normalized killing threshold 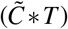 for models using a constant sink strength *a*_*s*_ (**a**), a metabolic cost-adjusted sink strength *a*_*s*_(1 − *µ/µ* * WT) (**b**), or WT architecture combined with metabolic cost adjustment (**c**). **d**, Scatter plots comparing predicted bulk survival with experimental values for various sink strengths (*a*_*s*_) across different modeling assumptions: constant (*C*), metabolic cost-adjusted (*M*), or WT architecture combined with metabolic cost adjustment (*M/WT*). **e–g**, Fitted heterogeneity *ℋ* as a function of the dimensionless relative screening length under constant *a*_*s*_ (**e**), metabolic cost-adjusted *a*_*s*_ (**f**), or WT architecture combined with metabolic cost adjustment (**g**).

**Supplementary Fig. 18:**
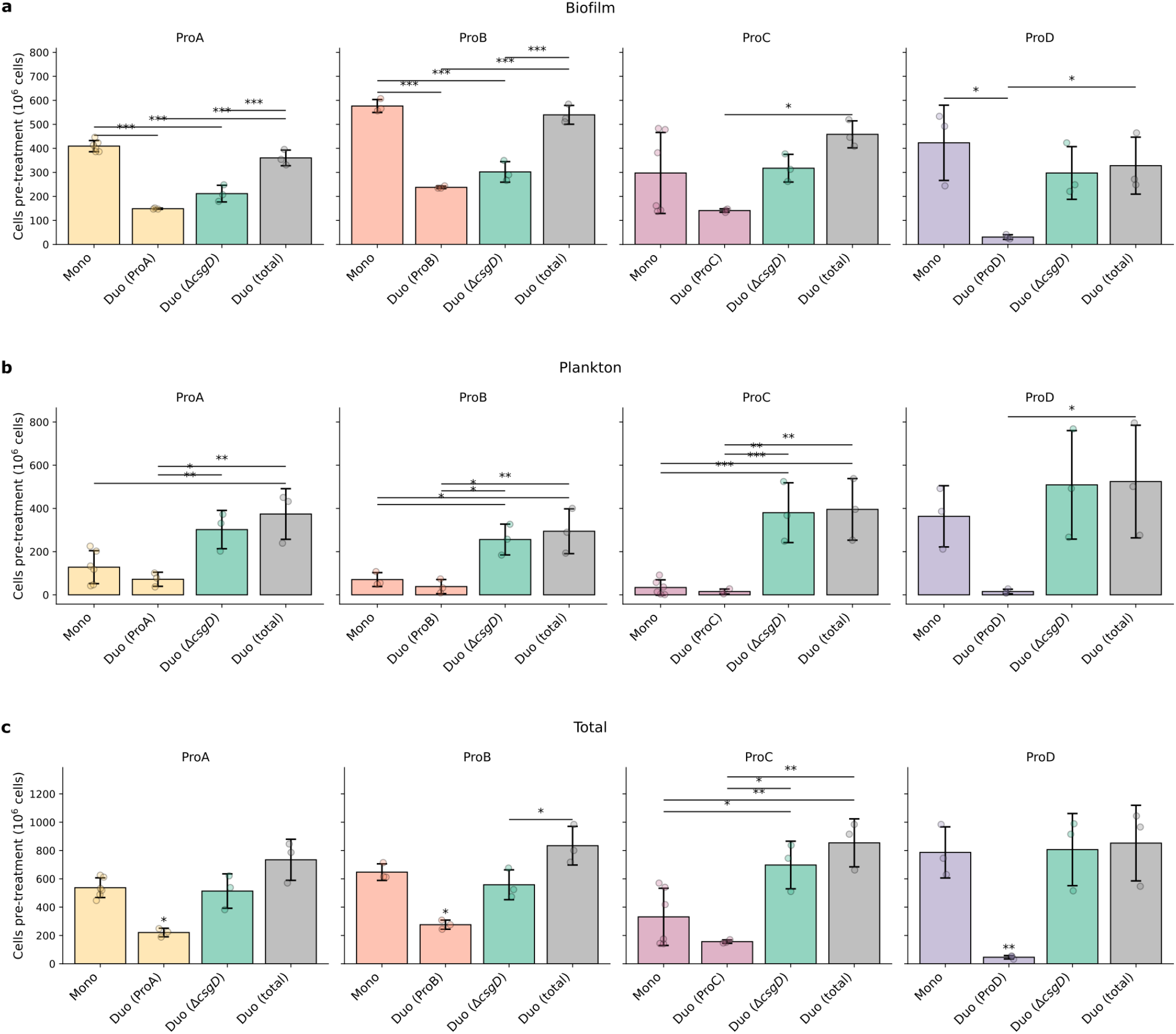
Co-culture of strains with engineered EPS levels and the EPS-deficient mutant Δ*csgD*. Comparison of the number of cells after 24 h of growth for the respective EPS-producing strain (ProA–D) in monoculture and in co-culture with a 1:1 inoculum of the EPS-deficient mutant Δ*csgD*. Cell counts for the monoculture, the respective strain within the co-culture, the Δ*csgD* mutant, and the overall co-culture are compared for the biofilm (**a**), planktonic (**b**), and combined (**c**) populations. Points indicate the average of three technical repeats for each of three independent biological repeats, whereas bars and error bars indicate the mean and standard deviation across the *n* = 3 biological repeats. Statistically significant pairwise comparisons were determined using one-way analysis of variance (ANOVA) followed by Tukey’s honestly significant difference (HSD) test. ^***^*P* < 0.001; ^**^*P* < 0.01; ^*^*P* < 0.05; n.s., not significant.

**Supplementary Fig. 19:**
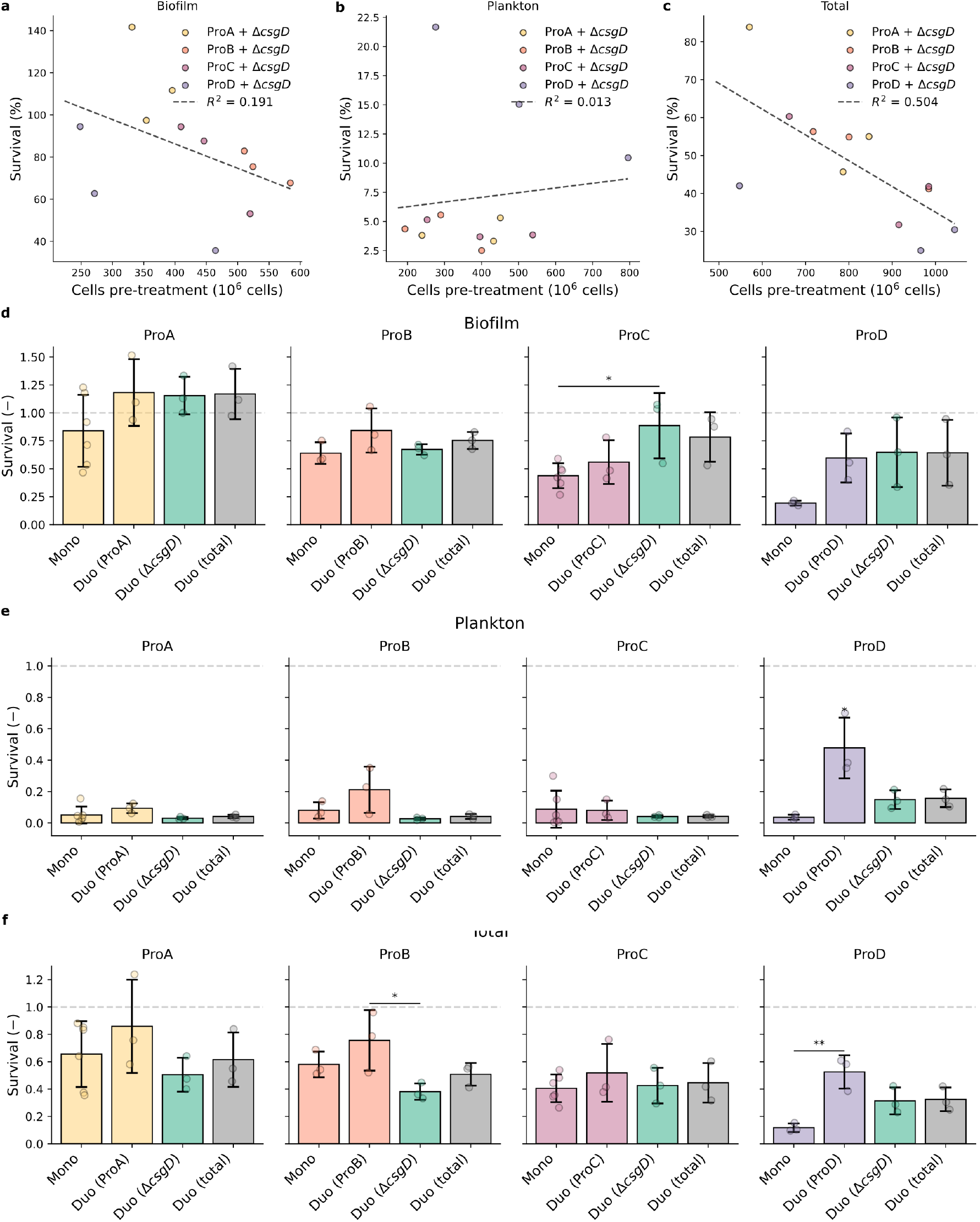
Bulk antimicrobial survival following cefotaxime treatment of strains co-cultured with the EPS-deficient mutant Δ*csgD*. **a–c**, No strong correlation was observed between the total number of cells within the co-cultured biofilms before treatment and eventual overall survival for the planktonic (**a**), biofilm (**b**), and total (**c**) populations. **d–f**, Differences in survival between monocultures of the respective EPS-producing strain and the survival of the respective strain in co-culture, the EPS-deficient mutant Δ*csgD* in co-culture, and the total survival of the co-culture for the planktonic (**d**), biofilm (**e**), and combined (**f**) populations. Points indicate the average of three technical repeats for each of three independent biological repeats, whereas bars and error bars indicate the mean and standard deviation across the *n* = 3 biological repeats. Statistically significant pairwise comparisons were determined using one-way analysis of variance (ANOVA) followed by Tukey’s honestly significant difference (HSD) test. ^***^*P* < 0.001; ^**^*P* < 0.01; ^*^*P* < 0.05; n.s., not significant.

## References

1. Costerton, J. W., Stewart, P. S. & Greenberg, E. P. Bacterial Biofilms: A Common Cause of Persistent Infections. Science 284, 1318–1322, DOI: 10.1126/science.284.5418.1318 (1999).

2. Flemming, H.-C. & Wingender, J. The biofilm matrix. Nat. Rev. Microbiol. 8, 623–633, DOI: 10.1038/nrmicro2415 (2010).

3. Ciofu, O., Moser, C., Jensen, P. O. & Høiby, N. Tolerance and resistance of microbial biofilms. Nat. Rev. Microbiol. 20, 621–635, DOI: 10.1038/s41579-022-00682-4 (2022).

4. Flemming, H.-C. et al. Biofilms: an emergent form of bacterial life. Nat. Rev. Microbiol. 14, 563–575, DOI: 10.1038/nrmicro.2016.94 (2016).

5. Stewart, P. S. Diffusion in Biofilms. J. Bacteriol. 185, 1485–1491, DOI: 10.1128/JB.185.5.1485-1491.2003 (2003).

6. Nadell, C. D., Foster, K. R. & Xavier, J. B. Emergence of Spatial Structure in Cell Groups and the Evolution of Cooperation. PLOS Comput. Biol. 6, e1000716, DOI: 10.1371/journal.pcbi.1000716 (2010).

7. Bravo, P., Lung Ng, S., MacGillivray, K. A., Hammer, B. K. & Yunker, P. J. Vertical growth dynamics of biofilms. Proc. Natl. Acad. Sci. 120, e2214211120, DOI: 10.1073/pnas.2214211120 (2023).

8. Stewart, P. S. et al. Reaction–diffusion theory explains hypoxia and heterogeneous growth within microbial biofilms associated with chronic infections. npj Biofilms Microbiomes 2, 16012, DOI: 10.1038/npjbiofilms.2016.12 (2016).

9. van Vliet, S., Hauert, C., Fridberg, K., Ackermann, M. & Dal Co, A. Global dynamics of microbial communities emerge from local interaction rules. PLOS Comput. Biol. 18, e1009877, DOI: 10.1371/journal.pcbi.1009877 (2022).

10. Dal Co, A., van Vliet, S., Kiviet, D. J., Schlegel, S. & Ackermann, M. Short-range interactions govern the dynamics and functions of microbial communities. Nat. Ecol. & Evol. 4, 366–375, DOI: 10.1038/s41559-019-1080-2 (2020).

11. van Gestel, J. et al. Short-range quorum sensing controls horizontal gene transfer at micron scale in bacterial communities. Nat. Commun. 12, 2324, DOI: 10.1038/s41467-021-22649-4 (2021).

12. Dal Co, A., van Vliet, S. & Ackermann, M. Emergent microscale gradients give rise to metabolic cross-feeding and antibiotic tolerance in clonal bacterial populations. Philos. Transactions Royal Soc. B: Biol. Sci. 374, 20190080, DOI: 10.1098/rstb.2019.0080 (2019).

13. Persat, A. et al. The mechanical world of bacteria. Cell 161, 988–997, DOI: 10.1016/j.cell.2015.05.005 (2015).

14. Hartmann, R. et al. Emergence of three-dimensional order and structure in growing biofilms. Nat. Phys. 15, 251–256, DOI: 10.1038/s41567-018-0356-9 (2019).

15. Jeckel, H. et al. Shared biophysical mechanisms determine early biofilm architecture development across different bacterial species. PLOS Biol. 20, e3001846, DOI: 10.1371/journal.pbio.3001846 (2022).

16. Qin, B. et al. Cell position fates and collective fountain flow in bacterial biofilms revealed by light-sheet microscopy. Science 369, 71–77, DOI: 10.1126/science.abb8501 (2020).

17. Nijjer, J. et al. Biofilms as self-shaping growing nematics. Nat. physics 19, 1936–1944, DOI: 10.1038/s41567-023-02221-1 (2023).

18. Beroz, F. et al. Verticalization of bacterial biofilms. Nat. Phys. 14, 954–960, DOI: 10.1038/s41567-018-0170-4 (2018).

19. Chirwa, N. T. & Herrington, M. B. CsgD, a regulator of curli and cellulose synthesis, also regulates serine hydroxymethyltransferase synthesis in Escherichia coli K-12. Microbiology 149, 525–535, DOI: 10.1099/mic.0.25841-0 (2003).

20. Zogaj, X., Bokranz, W., Nimtz, M. & Römling, U. Production of cellulose and curli fimbriae by members of the family Enterobacteriaceae isolated from the human gastrointestinal tract. Infect. Immun. 71, 4151–4158, DOI: 10.1128/IAI.71.7.4151-4158.2003 (2003).

21. Gerstel, U. & Römling, U. The csgD promoter, a control unit for biofilm formation in Salmonella typhimurium. Res. Microbiol. 154, 659–667, DOI: 10.1016/j.resmic.2003.08.005 (2003).

22. Barnhart, M. M. & Chapman, M. R. Curli biogenesis and function. Annu. Rev. Microbiol. 60, 131–147, DOI: 10.1146/annurev.micro.60.080805.142106 (2006).

23. Thongsomboon, W. et al. Phosphoethanolamine cellulose: A naturally produced chemically modified cellulose. Science 359, 334–338, DOI: 10.1126/science.aao4096 (2018).

24. Serra, D. O., Richter, A. M. & Hengge, R. Cellulose as an architectural element in spatially structured Escherichia coli biofilms. J. Bacteriol. 195, 5540–5554, DOI: 10.1128/JB.00946-13 (2013).

25. Davis, J. H., Rubin, A. J. & Sauer, R. T. Design, construction and characterization of a set of insulated bacterial promoters. Nucleic Acids Res. 39, 1131–1141, DOI: 10.1093/nar/gkq810 (2011).

26. Grantcharova, N., Peters, V., Monteiro, C., Zakikhany, K. & Römling, U. Bistable expression of CsgD in biofilm development of Salmonella enterica serovar typhimurium. J. Bacteriol. 192, 456–466, DOI: 10.1128/JB.01826-08 (2010).

27. van Bruggen, M. P. B., Dhont, J. K. G. & Lekkerkerker, H. N. W. Morphology and Kinetics of the Isotropic-Nematic Phase Transition in Dispersions of Hard Rods. Macromolecules 32, 2256–2264, DOI: 10.1021/ma981196e (1999).

28. Olsen, I. Biofilm-specific antibiotic tolerance and resistance. Eur. J. Clin. Microbiol. & Infect. Dis. 34, 877–886, DOI: 10.1007/s10096-015-2323-z (2015).

29. Drescher, K. et al. Architectural transitions in Vibrio cholerae biofilms at single-cell resolution. Proc. Natl. Acad. Sci. 113, E2066–E2072, DOI: 10.1073/pnas.1601702113 (2016).

30. Serra, D. O. & Hengge, R. A c-di-GMP-Based Switch Controls Local Heterogeneity of Extracellular Matrix Synthesis which Is Crucial for Integrity and Morphogenesis of Escherichia coli Macrocolony Biofilms. J. Mol. Biol. 431, 4775–4793, DOI: 10.1016/j.jmb.2019.04.001 (2019).

31. Moreau, A. et al. Surface remodeling and inversion of cell-matrix interactions underlie community recognition and dispersal in Vibrio cholerae biofilms. Nat. Commun. 16, 327, DOI: 10.1038/s41467-024-55602-2 (2025).

32. Cordisco, E., Zanor, M. I., Moreno, D. M. & Serra, D. O. Selective inhibition of the amyloid matrix of Escherichia coli biofilms by a bifunctional microbial metabolite. npj Biofilms Microbiomes 9, 81, DOI: 10.1038/s41522-023-00449-6 (2023).

33. Hancock, A. M. et al. A nutrient bottleneck controls antibiotic efficacy in structured bacterial populations. Nat. Commun. 17, 3337, DOI: 10.1038/s41467-026-69625-4 (2026).

34. Kumar, A. & Schweizer, H. P. Bacterial resistance to antibiotics: Active efflux and reduced uptake. Adv. Drug Deliv. Rev. 57, 1486–1513, DOI: 10.1016/j.addr.2005.04.004 (2005).

35. Denk-Lobnig, M. K. & Wood, K. B. Spatial population dynamics of bacterial colonies with social antibiotic resistance. Proc. Natl. Acad. Sci. 122, e2417065122, DOI: 10.1073/pnas.2417065122 (2025).

36. Yurtsev, E. A., Chao, H. X., Datta, M. S., Artemova, T. & Gore, J. Bacterial cheating drives the population dynamics of cooperative antibiotic resistance plasmids. Mol. Syst. Biol. 9, 683, DOI: 10.1038/msb.2013.39 (2013).

37. Datsenko, K. A. & Wanner, B. L. One-step inactivation of chromosomal genes in Escherichia coli K-12 using PCR products. Proc. Natl. Acad. Sci. United States Am. 97, 6640–6645, DOI: 10.1073/pnas.120163297 (2000).

38. Santiviago, C. A. et al. Analysis of Pools of Targeted Salmonella Deletion Mutants Identifies Novel Genes Affecting Fitness during Competitive Infection in Mice. PLOS Pathog. 5, e1000477, DOI: 10.1371/journal.ppat.1000477 (2009).

39. Zwietering, M. H., Jongenburger, I., Rombouts, F. M. & van ’t Riet, K. Modeling of the Bacterial Growth Curve. Appl. Environ. Microbiol. 56, 1875–1881, DOI: 10.1128/aem.56.6.1875-1881.1990 (1990).

40. Choong, F. X. et al. Real-time optotracing of curli and cellulose in live salmonella biofilms using luminescent oligothiophenes. npj Biofilms Microbiomes 2, 1–11, DOI: 10.1038/npjbiofilms.2016.24 (2016). Number: 1 Publisher: Nature Publishing Group.

41. Frangi, A. F., Niessen, W. J., Vincken, K. L. & Viergever, M. A. Multiscale vessel enhancement filtering. In Wells, W. M., Colchester, A. & Delp, S. (eds.) Medical Image Computing and Computer-Assisted Intervention — MICCAI’98, 130–137, DOI: 10.1007/BFb0056195 (Springer, Berlin, Heidelberg, 1998).

42. Otsu, N. A Threshold Selection Method from Gray-Level Histograms. IEEE Transactions on Syst. Man, Cybern. 9, 62–66, DOI: 10.1109/TSMC.1979.4310076 (1979).

43. Hertz, H. Ueber die berührung fester elastischer körper. J. für die reine und angewandte Math. 93, 156–171 (1882).

44. Alexander, S. Adsorption of chain molecules with a polar head a scaling description. J. de Physique 38, 983–987, DOI: 10.1051/jphys:01977003808098300 (1977).

45. de Gennes, P. G. Polymers at an interface; a simplified view. Adv. Colloid Interface Sci. 27, 189–209, DOI: 10.1016/0001-8686(87)85003-0 (1987).

46. Cooper, K. G., Chong, A., Starr, T., Finn, C. E. & Steele-Mortimer, O. Predictable, Tunable Protein Production in Salmonella for Studying Host-Pathogen Interactions. Front. Cell. Infect. Microbiol. 7, DOI: 10.3389/fcimb.2017.00475 (2017).

47. Kim, J., Webb, A. M., Kershner, J. P., Blaskowski, S. & Copley, S. D. A versatile and highly efficient method for scarless genome editing in Escherichia coli and Salmonella enterica. BMC biotechnology 14, 84, DOI: 10.1186/1472-6750-14-84 (2014).

48. Mavridou, D. A. I., Gonzalez, D., Clements, A. & Foster, K. R. The pUltra plasmid series: A robust and flexible tool for fluorescent labeling of Enterobacteria. Plasmid 87-88, 65–71, DOI: 10.1016/j.plasmid.2016.09.005 (2016).

49. Zaslaver, A. et al. A comprehensive library of fluorescent transcriptional reporters for Escherichia coli. Nat. Methods 3, 623–628, DOI: 10.1038/nmeth895 (2006).

50. Andino, A. & Hanning, I. Salmonella enterica: Survival, Colonization, and Virulence Differences among Serovars. The Sci. World J. 2015, 520179, DOI: 10.1155/2015/520179 (2015).

51. Tuson, H. H. et al. Measuring the stiffness of bacterial cells from growth rates in hydrogels of tunable elasticity. Mol. Microbiol. 84, 874–891, DOI: 10.1111/j.1365-2958.2012.08063.x (2012).

52. Safari, A., Tukovic, Z., Walter, M., Casey, E. & Ivankovic, A. Mechanical properties of a mature biofilm from a wastewater system: from microscale to macroscale level. Biofouling 31, 651–664, DOI: 10.1080/08927014.2015.1075981 (2015).

53. Kundukad, B. et al. Mechanical properties of the superficial biofilm layer determine the architecture of biofilms. Soft Matter 12, 5718–5726, DOI: 10.1039/c6sm00687f (2016).

54. Sankaran, J. et al. Single microcolony diffusion analysis in Pseudomonas aeruginosa biofilms. npj Biofilms Microbiomes 5, 35, DOI: 10.1038/s41522-019-0107-4 (2019).

